# Structures of the Human Spliceosomes Before and After Release of the Ligated Exon

**DOI:** 10.1101/496695

**Authors:** Xiaofeng Zhang, Xiechao Zhan, Chuangye Yan, Wenyu Zhang, Dongliang Liu, Jianlin Lei, Yigong Shi

**Affiliations:** Beijing Advanced Innovation Center for Structural Biology, Tsinghua-Peking Joint Center for Life Sciences, School of Life Sciences and School of Medicine, Tsinghua University, Beijing 100084, China; Technology Center for Protein Sciences, Ministry of Education Key Laboratory of Protein Sciences, School of Life Sciences, Tsinghua University, Beijing 100084, China; School of Life Sciences, Westlake University, 18 Shilongshan Road, Xihu District, Hangzhou 310024, Zhejiang Province, China

## Abstract

Pre-mRNA splicing is executed by the spliceosome, which has eight major functional states each with distinct composition. Five of these eight human spliceosomal complexes, all preceding exon ligation, have been structurally characterized. In this study, we report the cryo-electron microscopy structures of the human post-catalytic spliceosome (P complex) and intron lariat spliceosome (ILS) at average resolutions of 3.0 and 2.9 Å, respectively. In the P complex, the ligated exon remains anchored to loop I of U5 small nuclear RNA, and the 3’-splice site is recognized by the junction between the 5’-splice site and the branch point sequence. The ATPase/helicase Prp22, along with the ligated exon and eight other proteins, are dissociated in the P-to-ILS transition. Intriguingly, the ILS complex exists in two distinct conformations, one with the ATPase/helicase Prp43 and one without. Comparison of these three late-stage human spliceosomes reveals mechanistic insights into exon release and spliceosome disassembly.

---

Pre-mRNA splicing, executed by the spliceosome, proceeds in two sequential steps: branching and exon ligation^1,2^. In the branching reaction, the 2’-OH from an invariant adenine nucleotide of the branch point sequence (BPS) serves as the nucleophile to attack the 5’-end phosphate of the 5’-splice site (5’SS), generating a free 5’-exon and an intron lariat-3’-exon intermediate^3^. In exon ligation, the newly exposed 3’-OH of the 5’-exon attacks the 5’-end phosphate of the 3’-exon, releasing the intron lariat and ligating the 5’-exon with the 3’-exon^3^. The fully assembled spliceosome exists in eight major functional states: precursor of the pre-catalytic spliceosome (pre-B), pre-catalytic spliceosome (B), activated spliceosome (B^act^), catalytically activated spliceosome (B^*^), catalytic step I complex (C), step II catalytically activated complex (C^*^), post-catalytic spliceosome (P), and intron lariat spliceosome (ILS)^4^. Branching and exon ligation occur in the B^*^ and C^*^ complexes, resulting in the C and P complexes, respectively.

Cryo-electron microscopy (cryo-EM) analysis of the spliceosome has revealed major insights into its assembly, activation, catalysis, and disassembly^5^. Through structural determination of all eight functional states of the spliceosome from *Saccharomyces cerevisiae* (*S. cerevisiae*)^1,6,7^, we have achieved mechanistic understanding of pre-mRNA splicing with unprecedented clarity. In contrast to the yeast studies, structural information of the human spliceosome has been slow to emerge^8-14^, largely due to its considerably more dynamic and unstable nature. In particular, the late-stage human spliceosomes (P and ILS) are yet to be structurally characterized.

In this manuscript, we report the atomic structures of the human P and ILS complexes at average resolutions of 3.0 and 2.9 Å, respectively. The ILS complex exists in two distinct conformations defined as ILS1 and ILS2. The ATPase/helicase Prp43, which is responsible for the disassembly of the spliceosome^15-17^, is only present in ILS2. Comparative analysis of these three late-stage human spliceosomes reveals the molecular mechanisms of 3’-splice site (3’SS) recognition, exon release and spliceosome disassembly.

## Spliceosome isolation and electron microscopy

The human spliceosomes were *in vitro* assembled on a synthetic MINX pre-mRNA. To enrich the late-stage spliceosomes, we purified an ATPase-defective mutant of Prp43 (with the missense mutation T429A)^18^ and assembled the spliceosome in the presence of an excess amount of the mutant Prp43 (Extended Data Fig. 1a,b). After affinity purification, the eluted sample was further fractionated by glycerol gradient centrifugation in the presence of the reagent glutaraldehyde. Chemical crosslinking ensures the integrity of the human spliceosomal particles^19^. The purified spliceosomes contained the intron lariat and a small amount of the ligated exon (Extended Data Fig. 1c). The particles appeared largely intact as revealed by EM under both negative-straining and cryogenic conditions (Extended Data Fig. 1d-f).

18,715 micrographs were recorded by a K2 Summit camera mounted on a Titan Krios microscope operating at 300 kV. Analysis of the first 2,000 micrographs revealed the presence of three major spliceosomal species (Extended Data Fig. 2a): one representing the P complex, to which the ligated exon remains bound, and the other two indicating the ILS complex, in which the exon is absent. A total of 2.1 million particles were auto-picked. Following three rounds of three-dimensional (3D) classification and subsequent refinement, 143,320 particles yielded a reconstruction of the human P complex at an average resolution of 3.0 Å (Fig. 1a; Extended Data Fig. 2b; Extended Data Tables 1 & 2). Using a similar strategy, we generated two distinct reconstructions of the human ILS complex. 390,072 and 499,840 particles gave rise to ILS1 and ILS2 at 2.91 and 2.86 Å, respectively (Fig. 1b,c; Extended Data Figs. 3 & 4; Extended Data Tables 1 & 3). The EM density map displays fine features for the bulk of the P complex (Extended Data Fig. 5) and the ILS complexes (Extended Data Fig. 6). Prp43 is present only in the ILS2 complex (Extended Data Fig. 6a).

**Figure 1.**
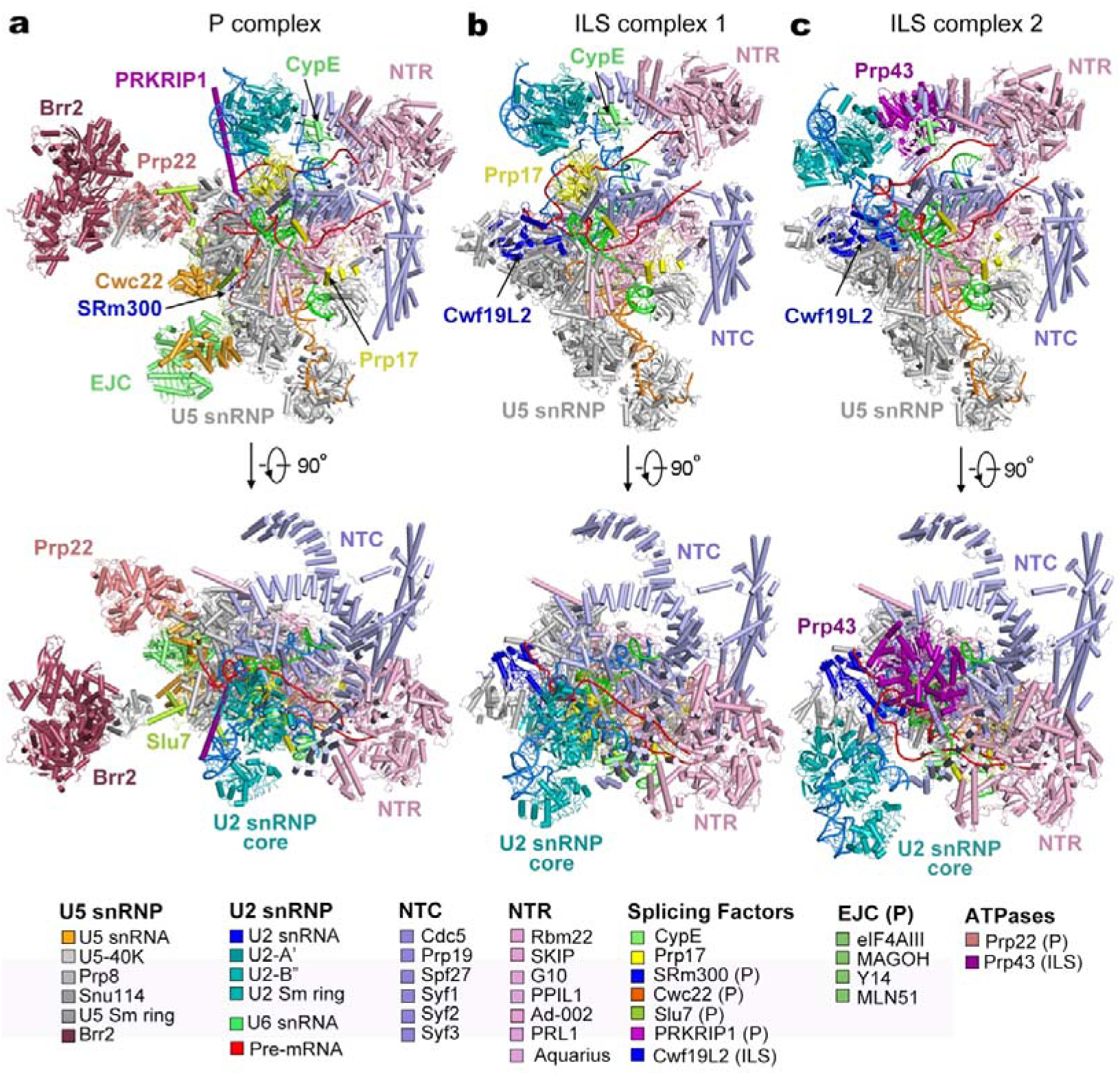
Cryo-EM structures of the human post-catalytic spliceosome (P complex) and the human intron lariat spliceosome (ILS complex). **a,** Structure of the human spliceosomal P complex at an average resolution of 3.0 Å. Two perpendicular views are shown. Components of the P complex are color-coded and tabulated below the structure. NTC: NineTeen complex; NTR: NTC related complex. **b,** Structure of the human ILS complex 1 (ILS1) at an average resolution of 2.91 Å. The ATPase/helicase Prp43 is yet to be loaded in the ILS1 complex. **c,** Structure of the human ILS complex 2 (ILS2) at an average resolution of 2.86 Å. Prp43 is present in the ILS2 complex, ready to disassemble the spliceosome.

## Overall structure of the human P complex

The atomic model of the human P complex contains 44 proteins and 4 RNAs, totaling 14,184 amino acids and 462 nucleotides (Extended Data Tables 1 & 2). The overall structure of the human P complex closely resembles that of the human C^*^ complex^12^ (Fig. 1a). In the P complex, the splicing factors SRm300 and Cwc22 and the exon junction complex (EJC) are bound to the upstream sequences of the 5’-exon (Fig. 1a). Portions of the step II splicing factors Prp17 and Slu7 adopt an extended conformation and interact with several proteins, stabilizing the active site conformation^20^. The extended α-helix of PRKRIP1 bridges the splicing active site with the core of U2 small nuclear protein (snRNP), with its N- and C-terminal portions interacting with the BPS/U2 duplex and the U2 small nuclear RNA (snRNA) duplex, respectively.

The pre-mRNA molecule consists of the intron lariat, which is stabilized by several protein components, and the ligated exon, which remains bound to loop I of U5 snRNA^21,22^. In the human C^*^ complex^12^, the 3’SS was not identified, presumably due to compromised recognition as a result of replacing the conserved dinucleotide AG by GG; In the human P complex, the 3’SS is clearly defined by the EM density map (Extended Data Fig. 5c). The ATPase/helicase Prp22, which is thought to dissociate the ligated exon by binding and pulling its 3’-end single-stranded RNA sequences^23,24^, is prominently anchored on the Linker domain of Prp8 (Fig. 1a). In the peripheral region of the P complex, Brr2 interacts with the Jab1/MPN domain of Prp8 (ref. 25) and the Brr2-Jab1 complex is connected to the spliceosomal core through interactions with the step II splicing factor Slu7 (ref. 26).

## Overall structure of the human ILS complex

In contrast to the P complex, the ILS complexes no longer contain the ligated exon (Fig. 1b,c). At least nine proteins have been dissociated in the P-to-ILS transition, presumably due to the action of Prp22. These proteins include four components of the EJC (eIF4AIII, MAGOH, MLN51, and Y14), the exon-stabilizing protein SRm300 (ref. 27), the splicing factors Cwc22 and Slu7, PRKRIP1, and the ATPase/helicase Prp22. Cwf19L1 and Cwf19L2 are the human homologs of *S. pombe* Cwf19^28^, with the former (Cwf19L1) known to be associated with recessive ataxia syndrome^29^. Notably, Cwf19L2, but not Cwf19L1, has been recruited into the ILS1 complex and forms extensive interactions with Prp8 and the intron lariat. Intriguingly, similar to the *S. pombe* ILS complex^30,31^, the ATPase/helicase Prp43 is yet to be loaded into the human ILS1 complex.

The only compositional difference between ILS1 and ILS2 is Prp43 (Fig. 1b,c). Similar to the *S. cerevisiae* ILS complex^32^, Prp43 is bound to the middle portion of the NTC component Syf1. Compared to those in the ILS1 complex, the core of U2 snRNP (U2-A’, U2-B’’ and the U2 Sm ring), the BPS/U2 duplex, and the splicing factor Prp17 have undergone pronounced movements (to be detailed later).

## Features of the human P complex

Exon ligation occurs spontaneously in the C^*^ complex, resulting in the P complex^3^. Therefore, the structures of the human P and C^*^ complexes should be nearly identical except the changes in pre-mRNA: the covalent linkage between the 3’SS and 3’-exon in the C^*^ complex is replaced by that between the 5’-exon and 3’-exon. Six contiguous nucleotides at the 3’-end of the intron, which are unambiguously identified in the EM density, are uniquely present in the human P complex (Fig. 2a). Recognition of the 3’SS is reminiscent of that in *S. cerevisiae*^33-35^. In particular, the guanine base of the 3’SS pairs up with the guanine (G1) at the 5’-end of the 5’SS through hydrogen bonds (H-bonds)^36,37^ and stacks against an adenine from nucleotide A45 of U6 snRNA (Fig. 2a, inset). A45, in turn, form two H-bonds with U2 of the 5’SS. The adenine base of the 3’SS, on the other hand, pairs up with the invariant adenine of the BPS and stacks against G1 of the 5’SS^38,39^. These interactions strictly depend on formation of the 2’-5’ linkage between the invariant adenine of the BPS and G1 of the 5’SS, which occurs in the branching reaction. These structural features ensure the temporal order of branching and exon ligation.

**Figure 2.**
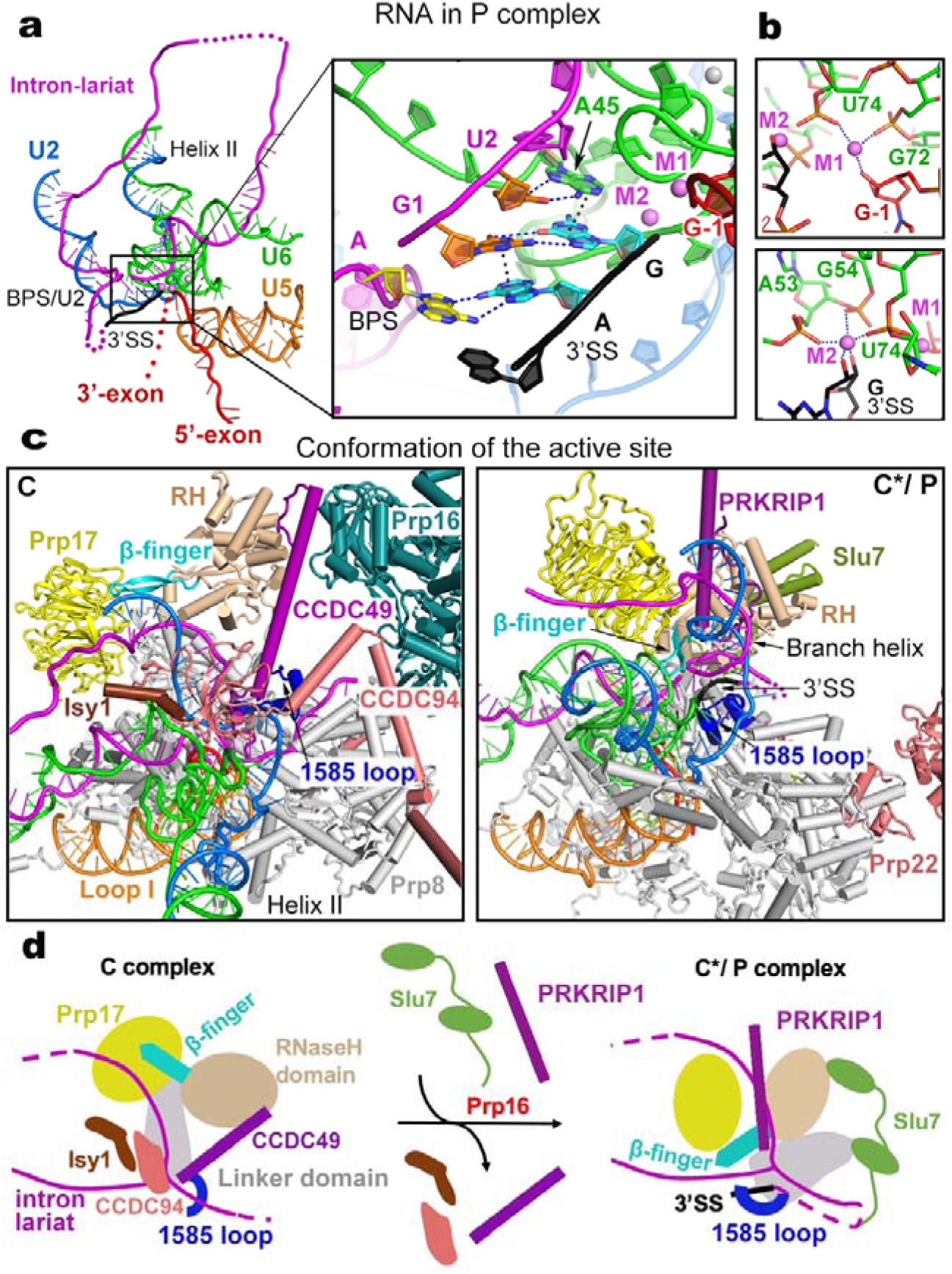
Structural features of the human P complex. **a,** Structure of the RNA elements in the P complex. An overall cartoon representation of the RNA map is shown on the left panel, and a close-up view on the splicing active site center is displayed in the right panel. Disordered RNA sequences in the intron lariat and the 3’-exon are represented by magenta and red dotted lines, respectively. The catalytic and structural metals are shown in magenta and gray spheres, respectively. The 3’-splice site (3’SS) is highlighted in black. Putative hydrogen bonds and van der Waals interactions that mediate recognition of the 3’SS are represented by blue and black dotted lines, respectively. **b,** Specific coordination of the catalytic metal ions in the human P complex. **c,** Structural rearrangements of the spliceosomal components at the active site during the C-to-P transition. The human C complex (left panel) and P complex (right panel) are shown in the same orientation as determined by the core of Prp8 and U5 snRNA. Shown here are the proteins and RNA elements around the catalytic center. **d,** A cartoon diagram of the 3’SS recognition. Remodeling of the C complex by Prp16 results in dissociation of the NTC component Isy1 and the step I factors CCDC49 and CCDC94/YJU2. The Linker domain of Prp8 is rotated away from the U2/BPS duplex. Consequently, the 1585-loop which binds the 3’-tail of the intron loads the 3’SS into the splicing active site center. The RNaseH-like domain is translocated towards the active site, pushing the β-finger towards the lariat junction. The WD40 domain of Prp17 is also translocated toward the branch helix; together with the splicing factor PRKRIP1 stabilize the new conformation of RNaseH-like domain and branch helix. The intron lariat junction is sandwiched by the β-finger and the 1585-loop. Slu7 adopts an extended conformation and binds the RNaseH-like and Linker domains of Prp8, stabilizing the local conformation.

Following exon ligation, the guanine nucleotide at the 3’SS has a freshly exposed 3’-OH that remains coordinated by the catalytic metal M2 (Fig. 2b). In addition, M2 is also coordinated by the 2’-OH of the same guanine and three phosphate oxygen atoms of the nucleotides A53, G54, and U74 of U6 snRNA. M1 is bound by two phosphate oxygens from G72 and U74 of U6 snRNA and the 3’-OH of G-1of the 5’-exon. Notably, The freshly exposed 3’-OH of the 3’SS is located about 5 Å away from the 3’-OH of the G-1 of the 5’-exon, consistent with the observation that exon ligation is reversible under certain conditions^40^.

In the published structure of the human C^*^ complex^12^, the catalytic center is incompletely defined due to the absence of the 3’SS and the moderate local resolution of 3.8 Å. In contrast, the 3’SS is specifically recognized in the catalytic center of the human P complex and the resolution of the local EM density map goes to 2.8 Å. We are now able to compare the catalytic centers between the human C complex and the P complex (which should faithfully reflect that of the C^*^ complex) (Fig. 2c). Compared to the C complex^10^, the NTC component Isy1 and the step I splicing factors CCDC49 (Cwc25 in *S. cerevisiae*) and CCDC94/YJU2 (Yju2 in *S. cerevisiae*) have been dissociated and the BPS/U2 duplex is rotated about 60 degrees in the C^*^/P complex (Fig. 2c,d). The WD40 domain of Prp17 and the RNaseH-like domain of Prp8 are translocated toward the space between the BPS/U2 and U6/5’SS duplexes, with the β-finger of the RNaseH-like domain inserted deeply into the active site of the C^*^/P complex. Placement of the β-finger into the active site is consistent with the observation that β-finger mutations rescue the exon ligation defects caused by mutations of the branch adenine and 3’SS^41^.

During the C-to-C^*^/P transition, PRKRIP1 is recruited and simultaneously interacts with the WD40 domain of Prp17, the RNaseH-like domain, and the BPS/U2 duplex. PRKRIP1 likely plays an important role in the C^*^/P complex by stabilizing the local conformation. The 1585-loop (also known as α-finger^42^) of the Linker domain undergoes conformational rearrangement to stabilize the placement of the 3’SS into the active site, with the pyrimidine-rich sequences between the BPS and 3’SS forming a loop in the open space (Fig. 2c, right panel); this loop is located within 20 Å of the modeled N-terminus of Slu7 (His64), consistent with the observation that Slu7 is involved in 3’SS selection^20,43,44^ and may play a role in the transfer of 3’SS to the active site^45^. In the human C^*^/P complex, the step II factor Slu7 also stabilizes the conformation of Prp8 through direct interactions with the RNaseH-like domain, endonuclease domain, Linker, and N-domain.

## The transition from the P to ILS1 complex

The transition from the P to the ILS complex is driven by Prp22, which presumably binds and pulls the 3’-end single-stranded RNA sequences of the ligated exon, resulting in its dissociation^23,24^. Compared to the human P complex, the overall organization of the human ILS1 complex remains largely unchanged, except that the ligated exon and at least nine protein components have been released (Fig. 3a; Extended Data Fig. 7a). The released proteins include the EJC, which binds the 5’-exon; SRm300, which stabilizes 5’-exon binding to loop I of U5 snRNA^27^; Cwc22, which stabilizes SRm300 and EJC^46-48^; PRKRIP1, which stabilizes the active site conformation; Slu7, which stabilizes Prp8; and Prp22, which triggers the P-to-ILS transition. With the departure of Slu7, the ATPase/helicase Brr2 and the Jab1 domain of Prp8 are only flexibly attached to the periphery of the spliceosome and thus no longer defined by the EM density map. Cwf19L2 is recruited to fill the space vacated by other proteins.

**Figure 3.**
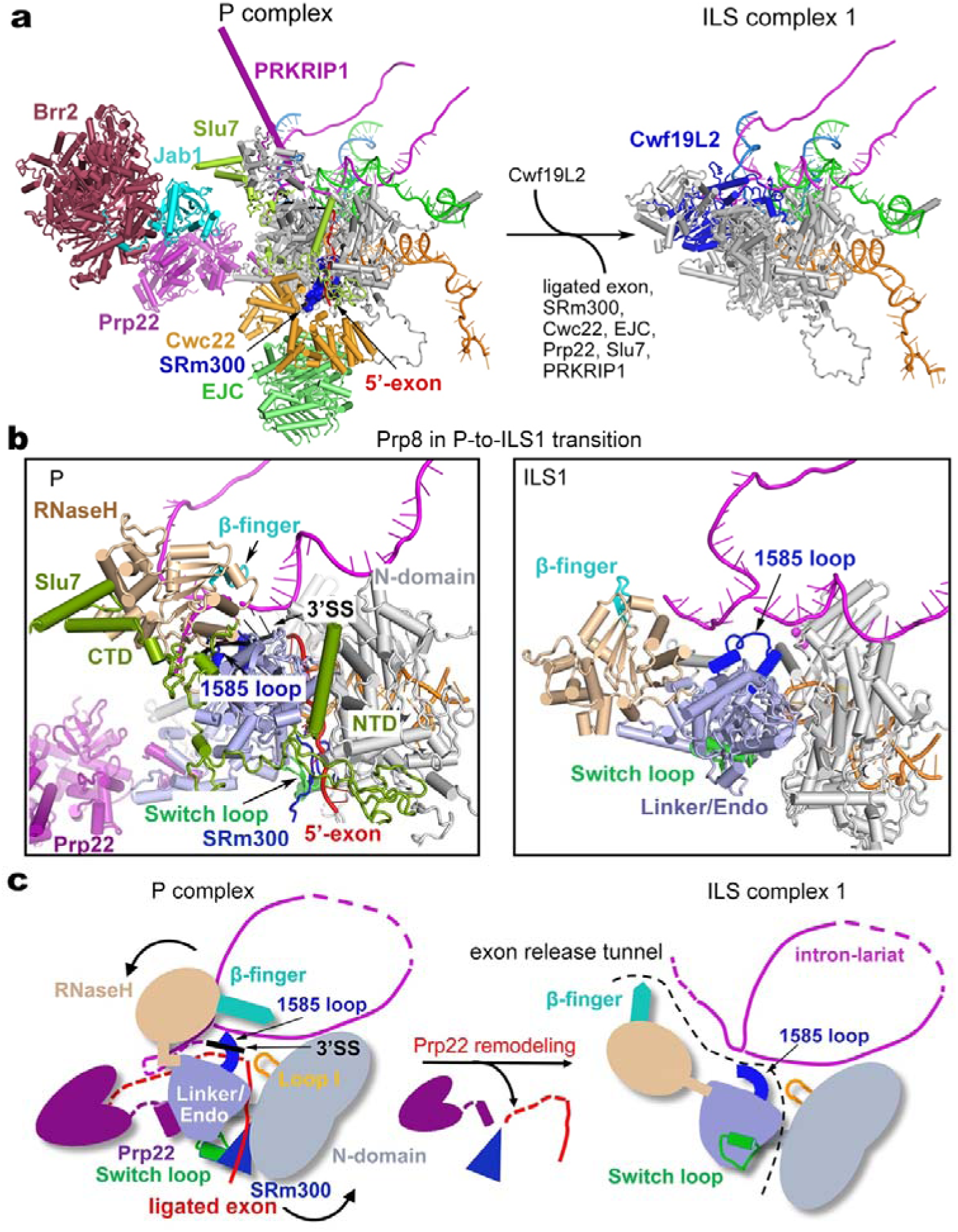
Structural rearrangements during the P-to-ILS1 transition. **a,** Overall structural changes during the P-to-ILS1 transition. All RNA elements are shown. The protein components that remain unchanged are not shown. SRm300 is shown in a surface representation. EJC: exon junction complex. **b,** Conformational changes of Prp8 during release of the ligated exon. During the P-to-ILS1 transition, the step II factor Slu7 and SRm300 is dissociated and the Switch loop that binds SRm300 is flipped by about 180 degrees. The RNaseH-like domain (colored wheat) and the 3’SS are moved away from the active site in the P complex. **c,** A cartoon diagram for the release of ligated exon. Driven by Prp22-mediated pulling, the β-finger (colored cyan) of the RNaseH-like domain is unbolted from the active site of the P complex; the Switch loop along with SRm300 and the 5’-exon are rotated 180 degrees, releasing the ligated exon through the tunnel formed by RNaseH-like, the Linker/Endo and N-domain of Prp8. In the P complex, the 1585-loop binds the 3’SS next to the exon junction, and this interaction must be disrupted prior to the release of the ligated exon. The 3’SS is flipped out of the catalytic center and becomes flexible during this process. The putative gate of exon release between intron lariat and four domains of Prp8 is marked by black dotted lines in the ILS1 complex.

Structural comparison of Prp8 and its surrounding components between the P and ILS1 complexes reveals mechanistic insights into exon release (Fig. 3b; Extended Data Fig. 8). In the P complex, the ligated exon is locked in the catalytic center formed by Prp8 and the RNA elements; these interactions are protected by surrounding protein components, particularly by Slu7. The Y-shaped N-terminal domain of Slu7 seals the potential exon-releasing cleft formed by the Linker, endonuclease domain, and N-domain of Prp8; the dumbbell-shaped C-terminal domain stabilizes the RNaseH-like domain (Fig. 3b, left panel). Driven by Prp22, the ligated exon is released in the P-to-ILS1 transition, causing efflux of the protein components and generating empty spaces for subsequent reorganization of the spliceosome. During the transition, the β-finger of the RNaseH-like domain is dislodged from, and rotated out of, the active site; the Switch loop, which binds SRm300, is flipped by about 180 degrees (Extended Data Fig. 8a). A small molecule, identified as inositol hexakisphosphate (IP6), remains bound to Prp8 during the P-to-ILS transition (Extended Data Fig. 8b). The 3’SS is released from the active site and becomes flexible again in the ILS1 complex.

Structural analysis reveals an exon-releasing tunnel formed by the intron lariat and four domains of Prp8 (N-domain, endonuclease, Linker, and RNaseH-like) in the P complex (Fig. 3b,c). We speculate that, driven by Prp22-mediated pulling from the 3’-end of the ligated exon, the step II factor Slu7 is released and the β-finger of the RNaseH-like domain is unbolted from the active site of the P complex. The Switch loop is flipped out by 180 degrees, allowing the release of SRm300 and threading of the exon first through the gate between the N-domain and endonuclease/Linker and then through the tunnel between the intron lariat and Prp8. Prior to exon release, the interactions between the 1585-loop and the 3’SS must be disrupted.

During the P-to-ILS1 transition, Cwf19L2 is recruited into the space between Prp8 and the intron lariat (Fig. 4a). Cwf19L2 closely interacts with the intron lariat and four domains of Prp8 (N-domain, endonuclease, Linker, and RNaseH-like), which contribute to formation of the putative exon release tunnel. These interactions stabilize the conformation of Prp8 and the intron lariat in the ILS1 complex. The final atomic model of Cwf19L2 (residues 536-894) comprises two domains linked by an extended bridging α-helix (Fig. 4b). Notably, a positively charged surface of the N-domain of Cwf19L2 clamps over the lariat junction(Fig. 4c), suggesting a role in modulating the turnover of branched intron^28^; whereas the bridging helix and the C-domain together bind the BPS/U2 duplex and the 3’-tail of the intron (Fig. 4c); this organization is consistent with the observation that Cwf19L2 may help displace and unwind the BPS/U2 duplex during spliceosome disassembly^49^.

**Figure 4.**
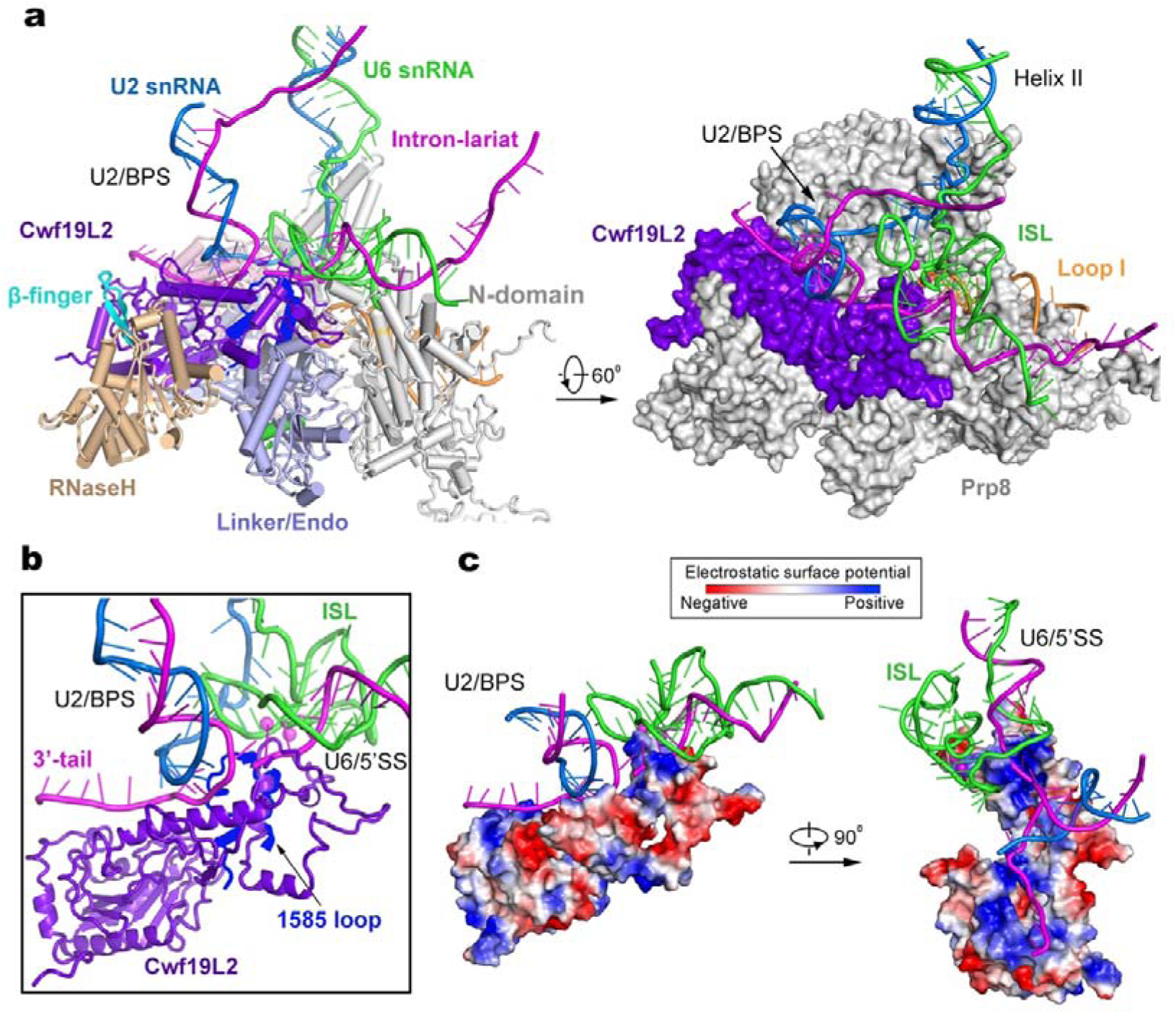
Cwf19L2 may play an important role during spliceosome disassembly and intron lariat RNA debranching. **a,** In the ILS1 complex, Cwf19L2 interacts closely with the lariat junction at the splicing active site and directly contacts the RNaseH-like (wheat), Linker/Endo (light blue), and N-domain (grey) of Prp8. Shown in the left is a cartoon representation of key components at the active site. Shown in the right is a surface representation of these components in a related view. **b,** A close-up view on the splicing active site. Prp8 is hidden except the 1585-loop (colored blue), which is located beneath the “clamp” of Cwf19L2. **c,** Cwf19L2 employs a positively charged surface to interact with the RNA elements. The electrostatic surface potential is shown for Cwf19L2. Two perpendicular views are presented.

## Recruitment of Prp43 and formation of the ILS2 complex

The only compositional difference between ILS1 and ILS2 is the ATPase/helicase Prp43 (Fig. 5a). Although the rigid core of the spliceosome remains unchanged between ILS1 and ILS2, binding by Prp43 results in marked positional shifts of surrounding components. Comparing to ILS1, the U2 snRNP core (U2-A’, U2-B’’ and the U2 Sm ring) undergoes a translocation in the ILS2 complex (Fig. 5b; Extended Data Fig. 6a; Extended Data Fig. 7b). The U2 snRNP core is displaced by about 30 Å away from Syf1. The helix IIc of U2 snRNA, which binds the RRM domain of CypE in ILS1, is co-translocated with the U2 snRNP core and consequently dissociated from CypE in ILS2. The RRM domain of CypE interacts with the intron sequence instead in the ILS2 complex. Intriguingly, the RRM domain of CypE binds single-stranded intron sequences in the human C complex^10^, switches over to interact with helix IIc of U2 snRNA in the C^*^/P/ILS1 complexes, and toggles back to bind the single-stranded intron sequences in the ILS2 complex (Extended Data Fig. 9). The PPI domain of CypE is only identified in the C complex. These structural features suggest a regulatory role for CypE in RNA splicing^50^. In the ILS2 complex, the U2 snRNP core becomes even more flexible (Extended Data Fig. 9), probably due to the initiation of spliceosome disassembly by the recruitment of Prp43. The constant positional shift of the U2 snRNP core is a hallmark of spliceosome remodeling during each splicing cycle^1^.

**Figure 5.**
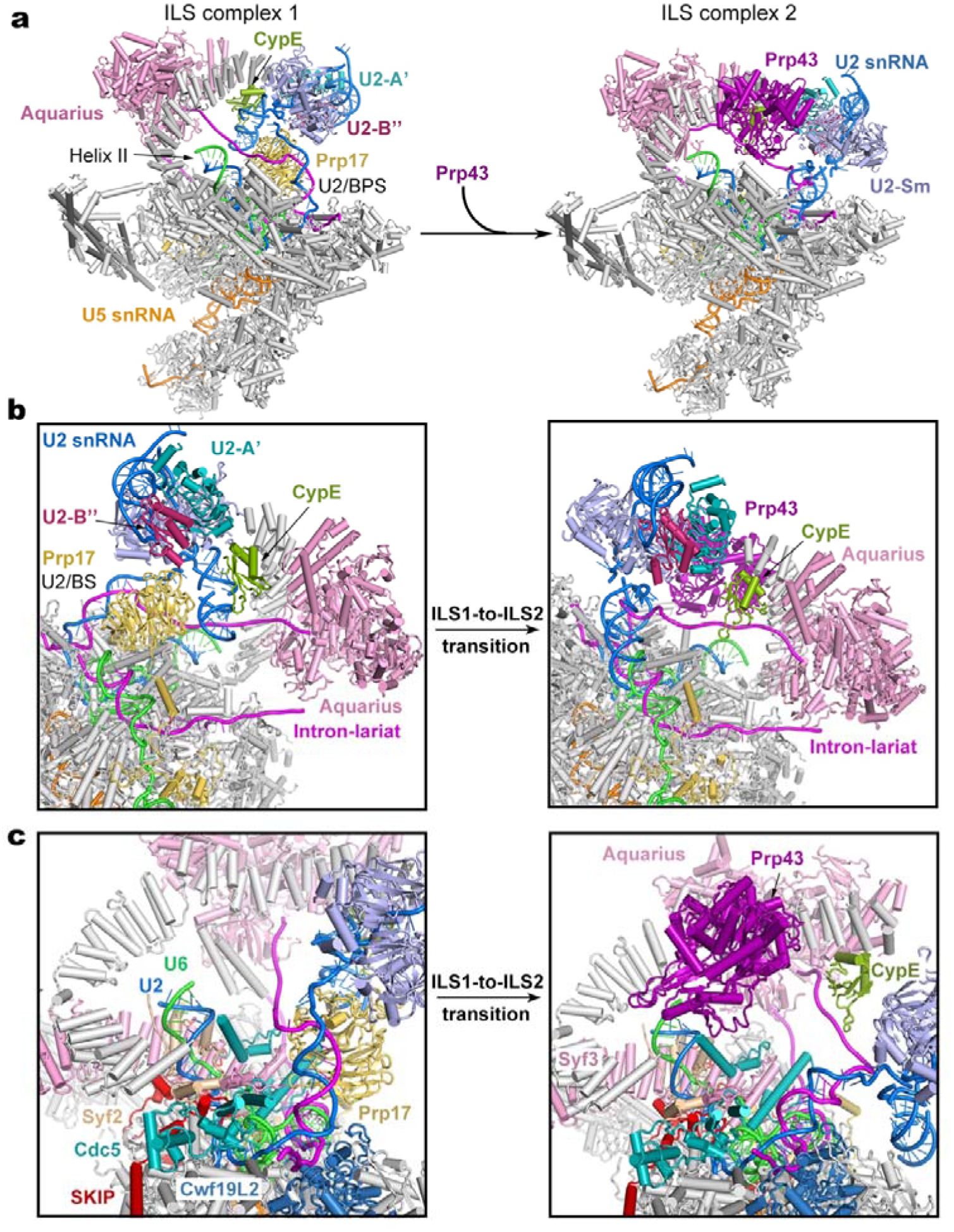
Structural rearrangements during the ILS1-to-ILS2 transition. **a,** Overall structural changes during the ILS1-to-ILS2 transition. The mobile protein components in the transition are color-coded to highlight the structural changes. Immobile proteins are shown in grey. In the transition, Prp43 is recruited into ILS2. **b,** A close-up view on select regions of the ILS1 and ILS2 complexes. Two orientations are shown. **c,** A close-up view on the transition from ILS1 to ILS2.

In the ILS1-to-ILS2 transition, the BPS/U2 duplex and the ensuing intron sequences are rotated by about 45 degrees (Fig. 5b). Consequently, the WD40 domain of Prp17, which binds the BPS/U2 duplex and U2 helix IIc in ILS1, is also translocated and becomes highly flexible and can no longer be identified in the EM density map. As previously observed in the *S. cerevisiae* ILS complex^32^, Prp43 is located in close proximity to helix II of the U2/U6 duplex (Fig. 5c). In the human ILS2 complex, Prp43 is positioned near the 3’-end of U6 snRNA, with a distance of about 40 Å between the RNA binding site of Prp43 and nucleotide 96 of U6 snRNA, consistent with extensive crosslinking between Prp43 and U6 snRNA^51^. This distance can be covered by seven consecutive nucleotides, allowing the 3’-end sequences to be grabbed and pulled by Prp43. Notably, we were able to trace the upstream sequences of BPS in the human ILS2 complex, but not in the *S. cerevisiae* ILS complex; this is likely due to the use of synthetic pre-mRNA for assembly of the human ILS2 complex. In fact, the intron sequences upstream of the BPS are sequentially bound by CypE and Aquarius and thus do not have sufficient space for Prp43 binding. These structural features are also consistent with the immunoprecipitation and crosslinking data that the intron sequence is not co-precipitated with, or crosslinked to, Prp43 (ref. 51). Lastly, the orientation of Prp43 also favors interaction with the 3’-end sequences of U6 snRNA instead of the intron sequence.

## Discussion

Using an ATPase-deficient Prp43 mutant, we were able to enrich human spliceosomes at late stages of the splicing cycle. The structures of the human P and ILS complexes fill the void for mechanistic understanding of pre-mRNA splicing by human spliceosome. Although the structure of the yeast P complex has been published^33-35^, improved resolution of the human P complex allows unambiguous identification of 3’SS recognition and catalytic metal coordination. The non-canonical base-pairing interactions between the BPS-5’SS junction and 3’SS is identical between the human and yeast P complexes. These observations confirm the suggested interactions between the first and last bases of the intron^52-54^ and explain the finding that mutations in the BPS led to deficiency of exon ligation^55-57^. Through comparison of the active site conformation in the C and C^*^/P complexes, we are able to propose a molecular mechanism for the placement and recognition of the 3’SS in the active site in the C-to-C^*^/P transition.

The human ILS complex has been uniquely captured in two distinct conformations. The ILS1 complex, in which Prp43 is yet to be recruited, may represent the conformational state right after Prp22 remolding. Following release of the ligated exon and its interacting proteins, CWF19L2 was recruited to occupy the vacated space in the active site, generating the ILS1 complex. Structural features of CWF19L2 support the idea that it likely plays a role in intron lariat RNA debranching^28^ and BPS/U2 translocation and spliceosome disassembly^49^. Through comparison between the P and ILS complexes, we identified a putative exon release tunnel, which is formed between intron lariat and four domains of Prp8.

Detailed structural information of the ILS complex is now available in *S. pombe*, *S. cerevisiae*, and human^30,32^ (Fig. 6a). Except U2 snRNP, all other conserved core components adopt nearly the same conformations in all three complexes.

**Figure 6:**
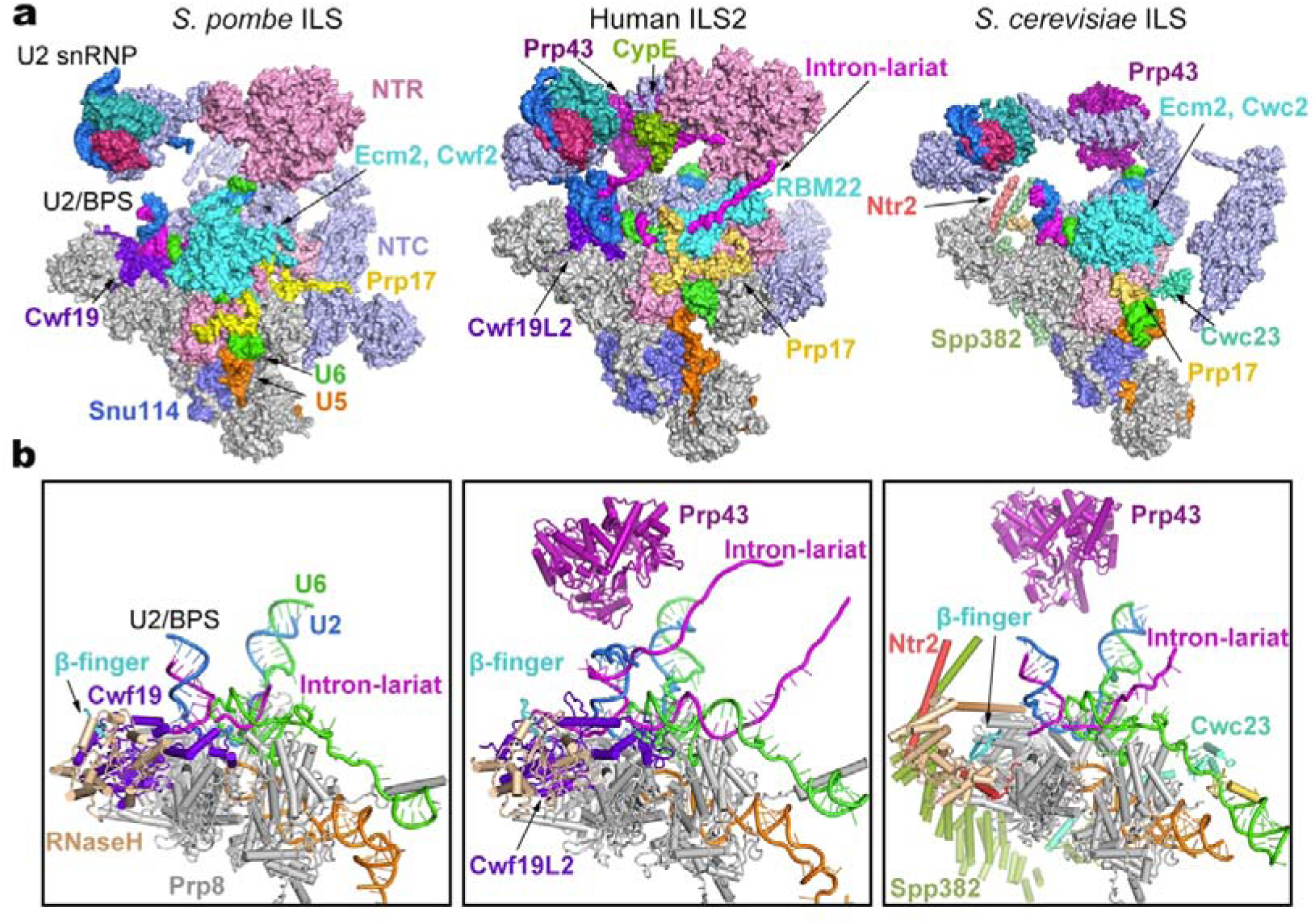
Structural comparison of the ILS complexes between yeast and human. **a,** Comparison of the overall structures of the ILS complex from *S. pombe* (left panel), human (middle panel), and *S. cerevisiae* (right panel). Components of the ILS complex are shown in surface representation. The components identically shared among the three complexes are shown in gray. Only those proteins that are different between the human and yeast complexes are color-coded. The *S. pombe* ILS complex resembles human ILS1; *S. pombe* Cwf19 (human Cwf19L2) closely interacts with Prp8 and the intron lariat, whereas Prp43 is yet to be loaded. The *S. cerevisiae* ILS lacks Cwf19 but contains 3 more proteins (Spp382/Ntr1, Ntr2, and Cwc23) that may participate in spliceosome disassembly. Notably, a large proportion of the RNA sequences in the intron lariat can be traced only in the human ILS complexes, but not in the yeast complexes. In the human ILS2 complex, the intron sequences upstream of the U2/BPS duplex is sequentially bound by CypE and Aquarius. The intron lariat traverses through a cavity formed by RBM22 in human. **b,** Comparison of the RNA elements and specific protein components in the ILS complex from *S. pombe* (left panel), human (middle panel), and *S. cerevisiae* (right panel). The RNaseH-like domain is colored wheat; the β-finger is marked by an arrow.

Notably, a large proportion of the RNA sequences in the intron lariat can only be traced in the human ILS complexes, but not in the yeast complexes. The human ILS1 complex closely resembles the *S. pombe* ILS complex, both lacking the ATPase/helicase Prp43 (Fig. 6b). In the human ILS2 complex, the intron sequences upstream of the U2/BPS duplex is sequentially bound by CypE and Aquarius. Another stretch of the intron lariat traverses through an internal cavity formed by RBM22 in human, instead of associating with Ecm2 and Cwc2/Cwf2 in yeast^58,59^.

The *S. pombe* ILS complex and the human ILS2 complex both contain a functionally mysterious protein Cwf19L2 (Cwf19 in *S. pombe*). In contrast to the ILS complex from *S. pombe* or human, the *S. cerevisiae* ILS complex is incapable of recruiting the Cwf19 homolog Drn1 (ref. 28). Instead, three proteins (Spp382/Ntr1, Ntr2 and Cwc23) are specifically recruited to facilitate the disassembly of the ILS complex^32^. The homologue of Ntr1 in *S. pombe* is yet to be identified. These observations suggest an evolutionary diversification of proteins involved in spliceosome disassembly. In addition, two lobes of unknown EM density are located close to Cwf19L2 and Prp43 in the human, but not yeast, ILS complexes (Extended Data Fig. 6a). These unidentified components might also play a role in the disassembly of human spliceosome.

The N-domain and the core of Prp8 are structurally superimposable in these three complexes. However, relative to the Prp8 core, the RNaseH-like domain adopts two different orientations, with the shared orientation between *S. pombe* and human different from that in *S. cerevisiae*. The distinct orientation of the RNaseH-like domain is likely facilitated by its close interactions with Cwf19L2 in human (Cwf19 in *S. pombe*). Our structural analysis and published studies support the conclusion that Cwf19L2 may have multiple roles during splicing^28^. The precise functional annotation of Cwf19L2 requires further biochemical investigation.

## Acknowledgments

We thank the Tsinghua University Branch of China National Center for Protein Sciences (Beijing) for the cryo-EM facility and the computational facility support, Xiaomin Li and Xiaofeng Hu for technical support in EM data acquisition. This work was supported by funds from the National Natural Science Foundation of China (31430020 and 31621092).

## Author Contributions

X. Zhang purified the human spliceosomes and prepared the cryo-EM samples. X. Zhang, X. Zhan, J.L. and C.Y. collected and processed the EM data. X. Zhang, X. Zhan, and C.Y. calculated the EM map. X. Zhan and C.Y. built the atomic model. All authors contributed to structure analysis and manuscript preparation. Y.S. designed and guided the project and wrote the manuscript.

## Author Information

The authors declare no competing financial interests. Correspondence and requests for materials should be addressed to Y.S.

## Methods

No statistical methods were used to predetermine sample size. The experiments were not randomized. The investigators were not blinded to allocation during experiments and outcome assessment.

### Expression and purification of human Prp43 mutants

Several dominant negative mutants of yeast Prp43 that abolish its NTPase or helicase activity have been previously identified^18^. The affected amino acids are evolutionally conserved from yeast to human. We individually introduced these mutations to human Prp43 (hPrp43) using a PCR-based strategy and subcloned the entire open reading frame into the expression vector pET21b with a C-terminal 6xHis tag. The resulting plasmid was transformed into *E. coli* BL21 codon plus strain. To increase the protein yield, ethanol was added to the cell culture to a final concentration of 3% (v/v) just prior to induction with IPTG^60^. The protein yield was increased up to 5 folds through the use of ethanol. Following Ni-NTA affinity purification, the eluted protein was further fractionated through heparin ion exchange chromatography to remove non-specifically bound nuclei acids. The peak fractions were combined and applied to gel filtration in buffer A (20mM HEPES-K 7.9, 450 mM KCl, 10% glycerol and 1 mM DTT). We used the purified Prp43 in the splicing assay and for analysis of the intron release efficiency. Among the several Prp43 mutants we tested, the mutation T429A abolished the ability of Prp43 to disassemble the ILS complex. This Prp43 mutant was chosen for subsequent experiments

### In vitro splicing reaction

The pre-mRNA sequence was modified from MINX and contains a 60-nucleotide (nt) 5’-exon, a 176-nt intron, and a 40-nt 3’-exon. The stem loop that contains two tandem MS2 binding sites is located in the middle of the intron. Based on structural information on human spliceosomes and prior biochemical characterization, such a design ensures that the binding of MS2-MBP protein does not interfere with the structural integrity of the spliceosome^12^. Pre-mRNA with a m7G(5’)ppp(5’)G cap was synthesized using the method of T7 runoff transcription. Splicing-active HeLa nuclear extract was prepared according to a published protocol^61^. A typical splicing reaction contains 25 nM pre-mRNA and 40% HeLa nuclear extract in a reaction buffer containing 2 mM ATP, 20 mM creatine phosphate, 20 mM HEPES–KOH 7.9, 65 mM KCl, and 3.5 mM MgCl_2_. To enrich the ILS complex, the dominant negative hPrp43 mutant (T429A) was pre-incubated with nuclear extract before initiation of the splicing reaction.

### Purification of the human ILS complex

We purified the human ILS complex using the strategy of MS2-MBP affinity purification essentially as previously described^12,62,63^. Briefly, pre-mRNA at a final concentration of 25 nM was incubated with MS2-MBP fusion protein at a final concentration of 500 nM on ice for 30 minutes. HeLa nuclear extract was supplemented with the hPrp43 (T429A) mutant protein to a final concentration of 100 ng/μl at room temperature for 30 minutes. Splicing reaction was initiated by adding splicing reagents and the pre-mRNA/MS2-MBP complex to the nuclear extract. The reaction was allowed to proceed at 30°C for 60 minutes. Early spliceosomal complexes and pre-mRNA that had not been incorporated into the spliceosome were digested by endogenous RNaseH with the addition of DNA oligonucleotides that are complementary to the upstream sequence of the 5’-splicing site (5’SS)^12^. The reaction was then loaded onto an amylose column for affinity purification. After extensive washing using the HS150 buffer (20 mM HEPES-KOH, 150 mM NaCl, 1.5 mM MgCl_2_, and 4% glycerol), the spliceosomal complexes were eluted by the HS150 buffer supplemented with 20 mM maltose. Splicing reactions were performed in a volume of 150 ml to obtain enough spliceosomal complexes for cryogenic electron microscopy (cryo-EM) studies. The eluted spliceosomal complexes were subjected to glycerol gradient centrifugation in the presence of 0-0.15% EM-grade glutaraldehyde. RNA components from peak fractions were analyzed on 8% Urea-PAGE.

### EM specimen preparation and data acquisition

The purified spliceosomal complexes were concentrated to ~0.25mg/ml for EM specimen preparation. Briefly, an aliquot of 3-μl sample was applied to a glow-discharged copper grid coated with a home-made carbon film. After waiting for 1 minute, the grid was blotted and immersed in uranyl acetate (2%, w/v) for negative staining or rapidly plunged into liquid ethane cooled by liquid nitrogen for cryo-EM specimen. To examine the integrity of the spliceosomes, negatively stained sample was imaged on a Tecnai Spirit Bio TWIN microscope operating at 120 kV. Cryo-EM grids were prepared using Vitrobot Mark IV (FEI Company) operating at 8°C and 100% humidity.

Cryo-EM specimen were imaged on a 300-kV Titan Krios electron microscope (FEI Company) with a normal magnification of 105,000x. Movies were recorded by a Gatan K2 Summit detector (Gatan Company) equipped with a GIF Quantum energy filter (slit width 20 eV) at the super-resolution mode, with a pixel size of 0.669 Å. Each stack of 32 frames was exposed for 8 seconds, with a dose rate of ~5.2 counts/second/physical-pixel (~4.7 e^-^/second/Å^2^) for each frame. AutoEMation^64^ was used for the fully automated data collection with high efficiency (~1800 stacks/24 hours). A total of 18,751 micrographs were collected. All 32 frames in each stack were aligned and summed using the whole-image motion correction program MotionCor2 (ref. 65) and binned to a pixel size of 1.338 Å. The defocus value for each image varied from 0.6 to 1.6 μm and was determined by Gctf^66^.

### Preliminary image processing

An initial dataset of 2,000 micrographs were first collected and processed to confirm the sample quality. 199,302 particles were auto-picked using the deep-learning program DeepPicker^67^. Reference-free two-dimensional (2D) classification was performed using RELION2.0 (refs. 68, 69), generating 150 classes. Good classes were combined, yielding 107,567 particles as an input for subsequent three-dimensional (3D) classification. The EM map of the human C^*^ complex^12^ (EMD-6721) was low-pass filtered to 40 Å as the initial reference. A single reference 3D classification was performed using the particles binned to a pixel size of 5.352 Å, yielding the P complex and two conformations of the ILS complex (named ILS1 and ILS2, respectively) based the fine features of the EM maps (Extended Data Fig. 2a).

### Image processing and calculation

In total, 2,104,128 particles were auto-picked from 18,715 micrographs using DeepPicker^67^. Guided multi-reference classification was applied to the full dataset using RELION2.0 (refs. 68, 69). Details of this modified procedure were previously described^10^. The 3D volumes of the P, ILS1 and ILS2 complexes generated from the preliminary data analysis, along with three bad classes, were low-pass filtered to 40 Å and used as the initial references (Round 1) (Extended Data Figs. 2b & 3). To avoid losing good particles, we simultaneously performed three parallel runs of multi-reference 3D classification. After Round 1, particles that belong to the P and ILS complexes were separated and used as the input for subsequent local classification. In each case, good particles from three parallel runs were merged, and the duplicated particles were removed. 588,094 particles (27.9% of the original input) for the P complex (Extended Data Fig. 2b), 460,373 particles (21.9%) for the ILS1 complex and 686,765 particles (32.6%) for the ILS2 complex were selected (Extended Data Fig. 3).

For P complex, a second round of multi-reference local 3D classification (Round 2) was performed using the 2x binned particles (pixel size: 2.676 Å). Good classes were merged and duplicated particles were removed to yield 173,916 particles. These 173,916 particles gave rise to a reconstruction of the human P complex at an average resolution of 4.65 Å after auto-refinement using unbinned particles (pixel size: 1.338 Å). After that, an additional round (Round 3) of 3D classification was performed. The remaining 173,916 particles were classified with a soft mask on the core region of the P complex. The class containing 143,320 particles (82.4% of the input) yielded a reconstruction at an average resolution of 3.24 Å of human P complex. These particles were further refined using THUNDER^70^. After global and local search, the resolution was improved to 3.04 Å (Extended Data Fig. 2b & 4a; Extended Data Table 1).

For the ILS complex, a second round of multi-reference local 3D classification (Round 2) was performed for the particles selected for ILS1 or ILS2 complex, respectively (Extended Data Fig. 3). Particles from the good classes were combined and duplicated particles were removed to yield 460,275 particles (21.9%) for ILS1 and 563,600 particles (26.8%) for ILS2. These particles gave rise to a reconstruction of the human ILS1 and ILS2 complexes at an overall resolution of 4.2 Å and 4.0 Å, respectively. Then an additional round of 3D classification (Round 3) was performed. The remaining 460,275 particles for ILS1 and 563,600 particles for ILS2 were classified with a soft mask on the core region. The class containing 390,072 particles (84.7% of the input particles) for ILS1 and 499,840 particles (88.7% of the input particles) for ILS2 each yielded a reconstruction at an average resolution of 3.2 Å. These particles were further refined using the THUNDER^70^; after global and local search, the final resolutions were improved to 2.91 Å for ILS1 and 2.86 Å for ILS2 (Extended Data Fig. 3 & 4a; Extended Data Table 1).

In the 3.0-Å map of the P complex, the local resolution reaches 2.8 Å in the core region of the spliceosome; in the 2.9-Å maps of both the ILS complexes, the local resolution reaches 2.8 Å or even higher (Extended Data Fig. 4b). The angular distributions of the particles used for the final reconstruction of human P and ILS complexes are reasonable (Extended Data Fig. 4c), and the refinement of the atomic coordinates did not suffer from severe over-fitting (Extended Data Fig. 4d). The resulting EM density maps display clear features for amino acid side chains in the core region of the human P and ILS complexes. The RNA elements and their interacting proteins are also well defined by the EM density maps.

Reported resolution limits were calculated on the basis of the FSC 0.143 criterion^71^, and the FSC curves were corrected with high-resolution noise substitution methods^72^. Prior to visualization, all EM maps were corrected for modulation transfer function (MTF) of the detector, and then sharpened by applying a negative B-factor that was estimated using automated procedures^71^. Local resolution variations were estimated using ResMap^73^.

### Model building and refinement

We combined de-novo model building, homologous structure modeling and rigid docking of components with known structures to generate the atomic models (Extended Data Tables 2 & 3). Identification and docking of the components of the human P and ILS complexes were facilitated by the structures of human C and C^*^ complexes^10,12^ and yeast ILS complex^30,32^. Protein components derived from known structures are summarized in Extended Data Tables S2 & S3. These structures were docked into the density map using Coot^74^ and fitted into density using CHIMERA^75^.

Briefly, the atomic coordinates of human C^*^ complex^12^ were directly docked into the 3.0-Å density map of the human P complex. The nucleotides at the 3’ end of the intron lariat were manually built using Coot^74^. The model of the RRM domain of CypE was generated from the human C complex^10^. The amino acid side chains of the protein components and the bases of the RNA nucleotides in the P complex were manually adjusted. On the basis of the EM density maps, six metal ions were identified. These metal ions were placed in the generally identical locations as those in the human C^*^ complex^12^. For both ILS complexes, the atomic coordinates derived from the human C^*^ complex^12^ were fitted into the 2.9-Å EM density maps. The protein and RNA components that are absent in the ILS complexes were removed, which include pre-mRNA, SRm300, Cwc22, Slu7, PRKRIP1, Prp22, and the exon junction complex (EJC). The atomic model of Cwf19L2 was generated from Cwf19 of the *S. pombe* ILS complex by CHAINSAW^76^. The automated model rebuilding was performed using RosettaCM with the adjusted model as the template and guided by the experimental cryo-EM density map^77-79^. Then the hydrogen atoms of the resulting model were removed and the atomic model was manually improved using Coot^74^. Crystal structure of human Prp43 (ref. 80) was docked into the density map of the ILS2 complex.

The final models of the human P and ILS complexes were refined according to the high quality cryo-EM maps using REFMAC in reciprocal space^81^ and secondary structure restraints that were generated by ProSMART^82^. Overfitting of the model was monitored by refining the model in one of the two independent maps from the gold-standard refinement approach, and testing the refined model against the other^83^ (Extended Data Fig. 4d). The structures of the human P and ILS complexes were validated through examination of the Molprobity scores and statistics of the Ramachandran plots (Extended Data Table 1). Molprobity scores were calculated as described^84^.

### Data availability.

The cryo-EM maps of the human P, ILS1 and ILS2 complexes have been deposited in the Electron Microscopy Data Bank (EMDB) with the accession codes EMD-9645, EMD-9646 and EMD-9647, respectively. The atomic coordinates of the human P, ILS1 and ILS2 complexes have been deposited in the Protein Data Bank (PDB) under the accession code 6ICZ, 6ID0 and 6ID1, respectably.

**Extended Data Table 1:**
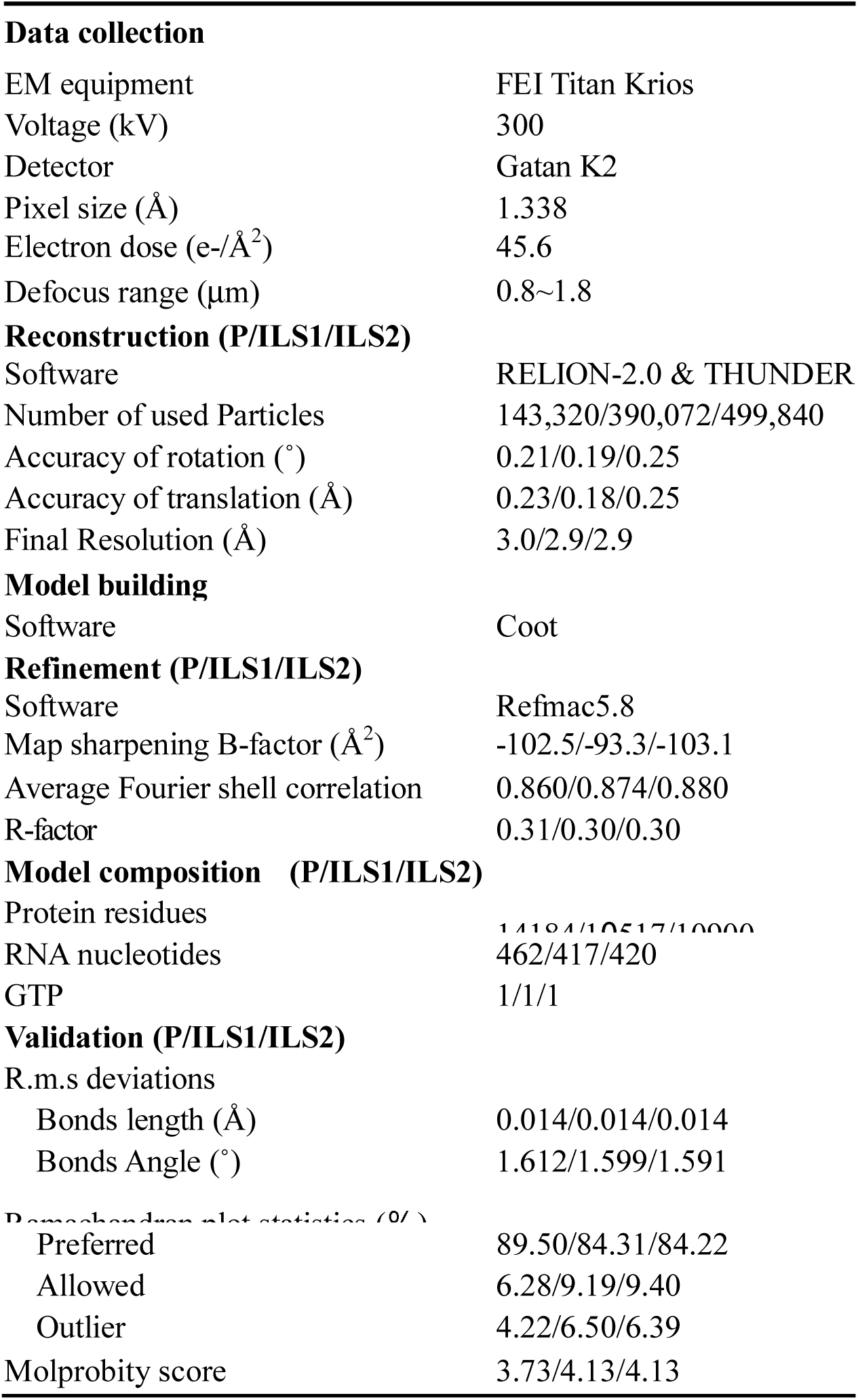
Cryo-EM data collection and refinement statistics.

**Extended Data Table 2:**
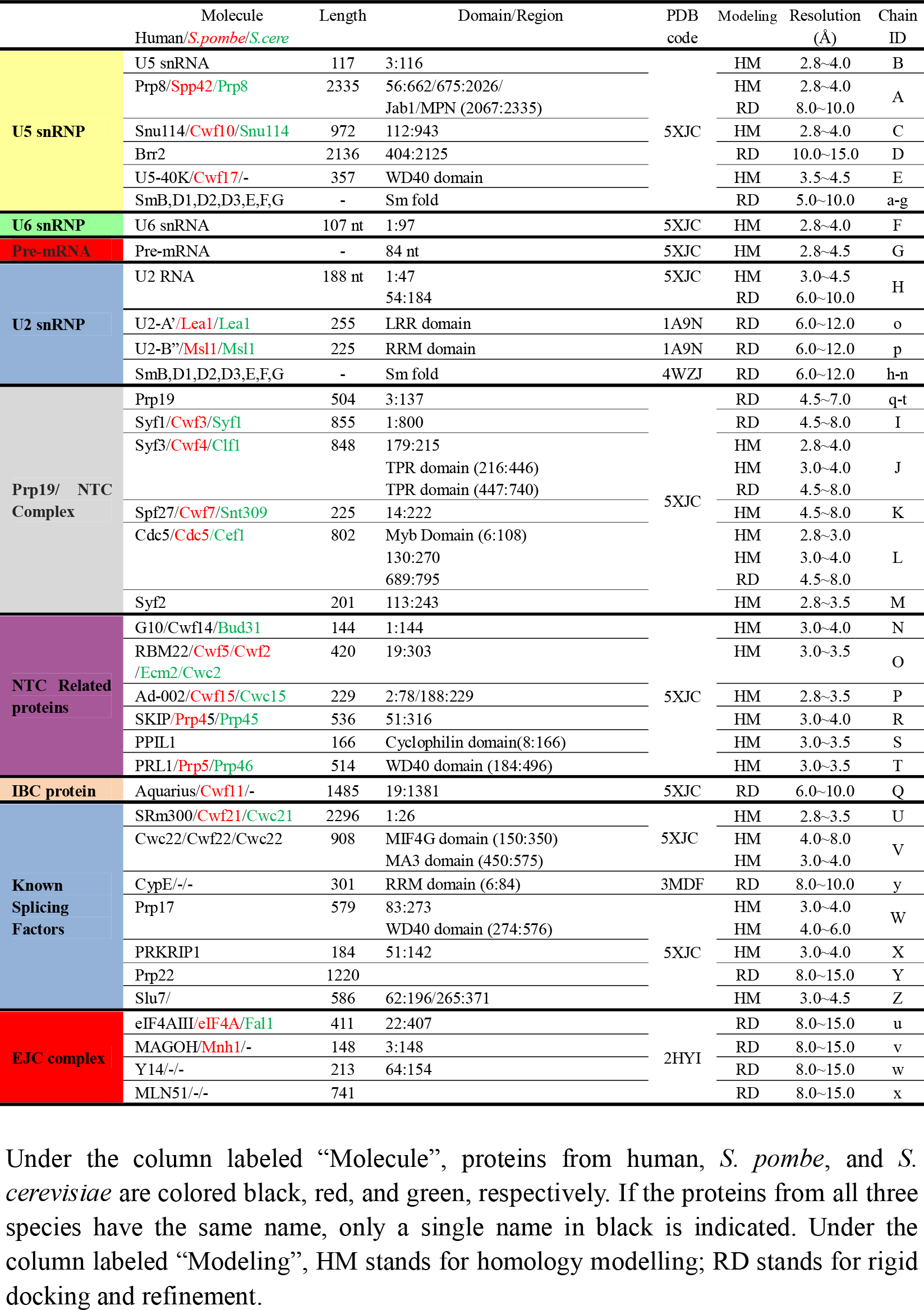
Summary of model building for the human spliceosomal P complex.

**Extended Data Table 3:**
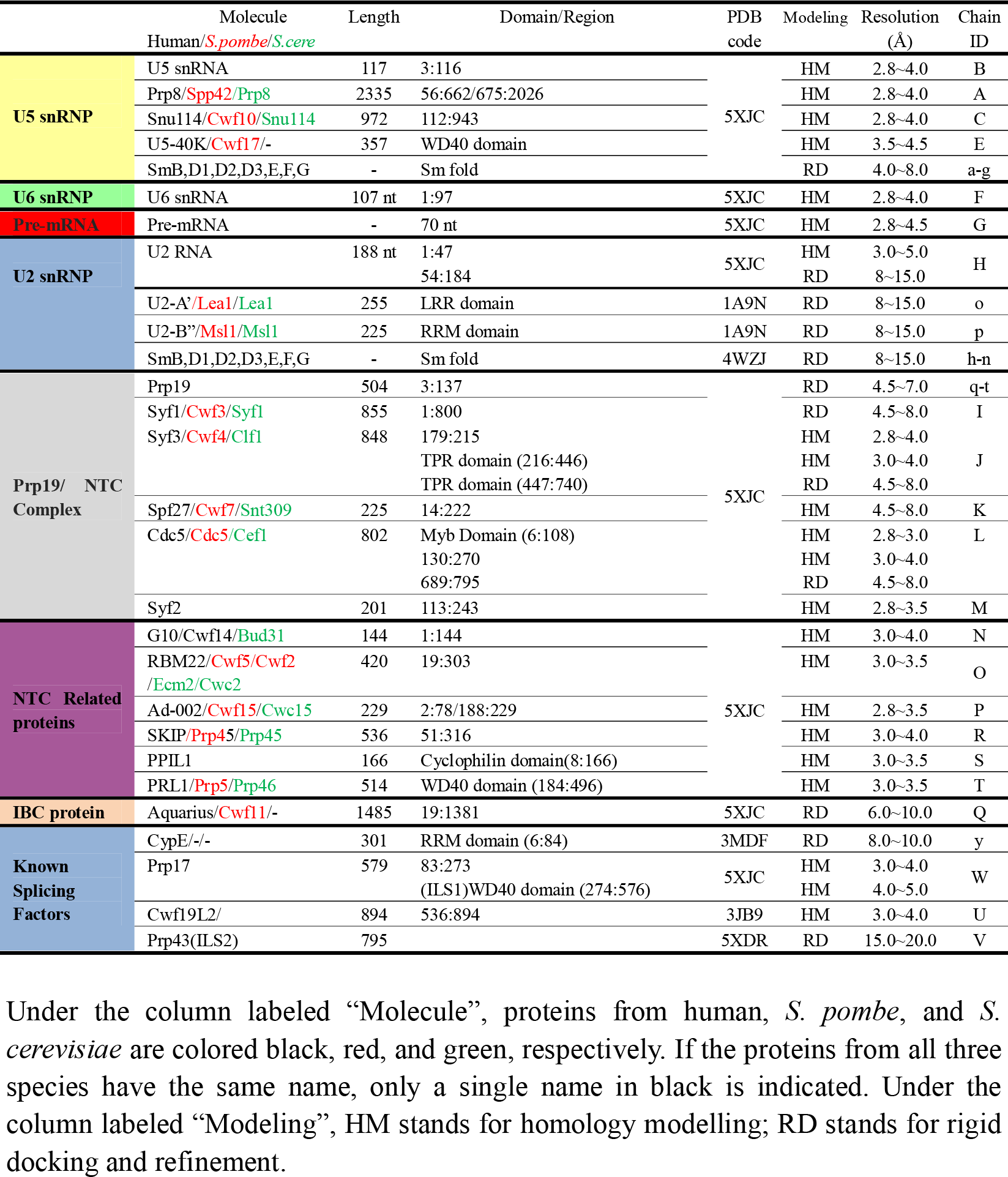
Summary of model building for the human spliceosomal ILS complex.

**Extended Data Figure 1:**
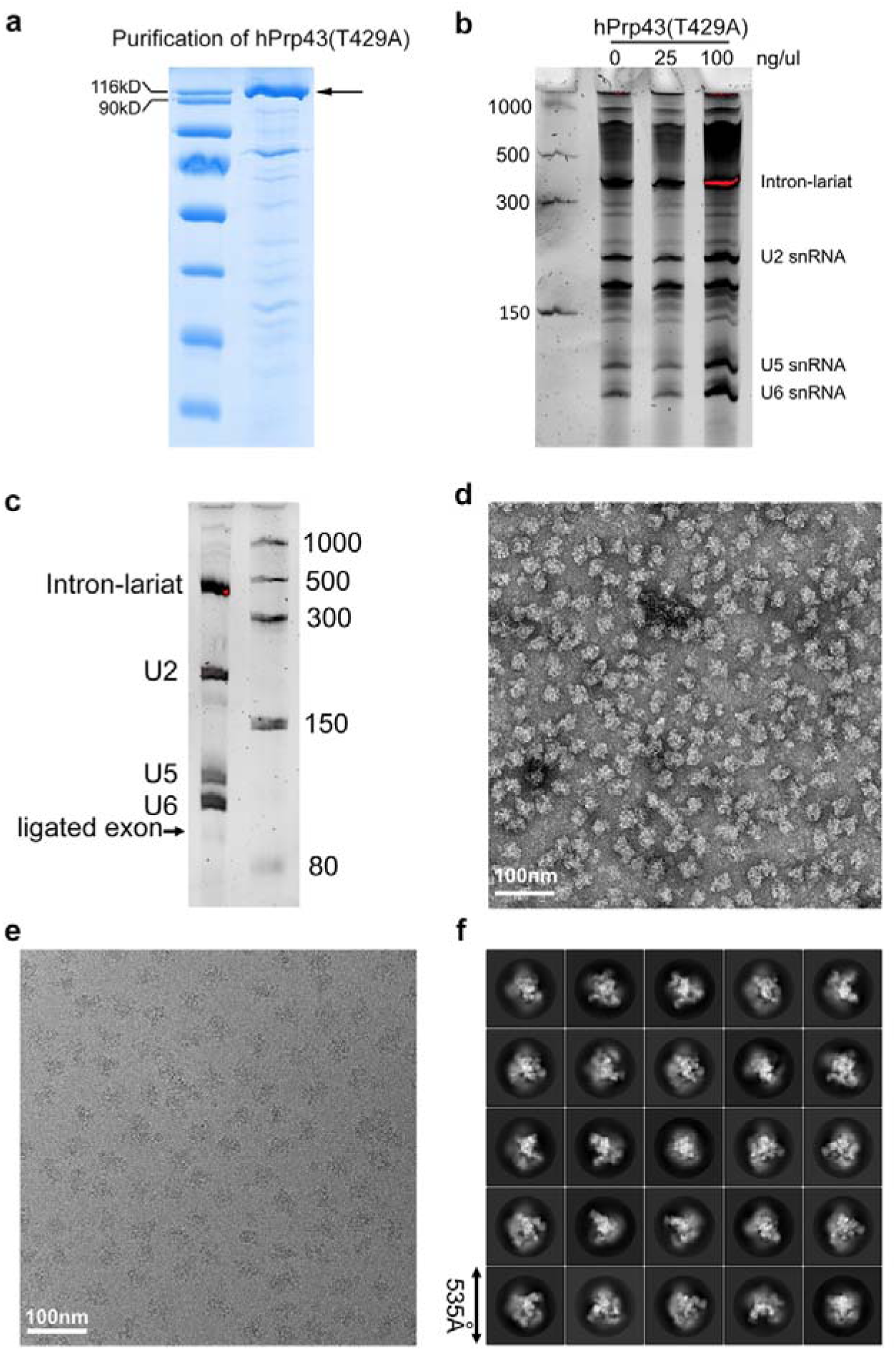
Purification and EM analysis of the human spliceosomal P and ILS complexes. **a**, The purified mutant protein Prp43 (T429A) is visualized by SDS-PAGE and Coomassie staining. **b,** Representative results of an intron release experiment shown on an RNA-urea gel. The splicing reaction was performed in the presence of varying amounts of hPrp43(T429A). The RNA elements of the ILS complex are indicated. **c,** An RNA gel showing the RNA components of the spliceosomal complexes after MS2-MBP affinity purification and glycerol gradient centrifugation. **d,** A representative electron microscopy (EM) micrograph of the final sample stained by uranyl acetate. Scale bar: 100 nm. **e,** A representative cryo-EM micrograph of the final sample. Scale bar: 100 nm. **f,** Representative 2D class averages calculated for the P and ILS complexes.

**Extended Data Figure 2:**
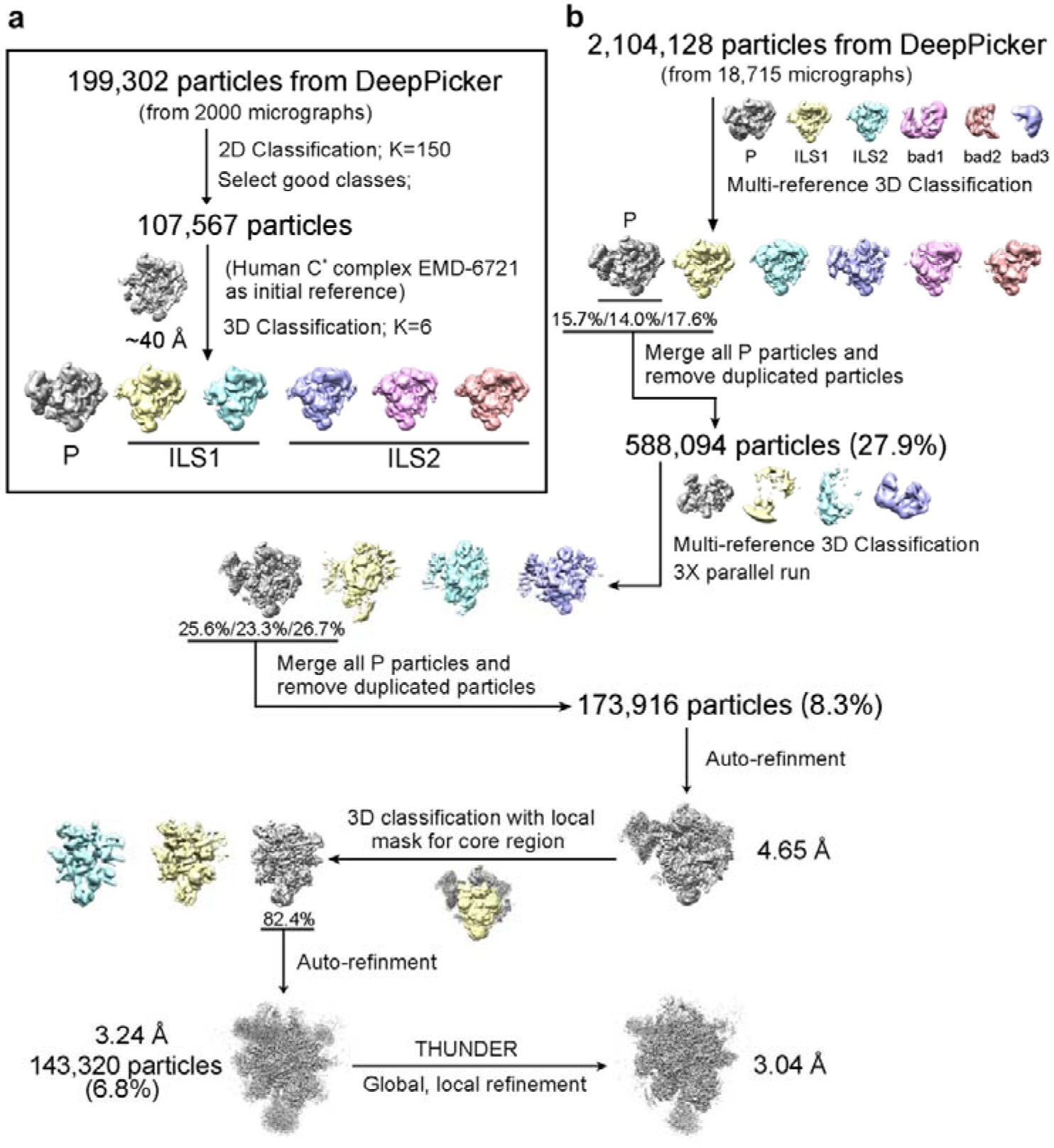
Preliminary cryo-EM data processing and structure determination of the human spliceosomal P complex. **a**, Analysis of the initial dataset of 2,000 micrographs of the affinity-purified human spliceosomal complexes. This analysis reveals that the vast majority of the particles belong to either the P complex (with ligated exon) or the ILS complex (ligated exon already dissociated). The ILS complex has two distinct conformations: one with Prp43 (ILS2) and one without Prp43 (ILS1). **b,** A flow chart description of the EM data processing and structure determination for the human spliceosomal P complex. The final reconstruction has an average resolution of 3.04 Å. Please refer to Methods for details.

**Extended Data Figure 3:**
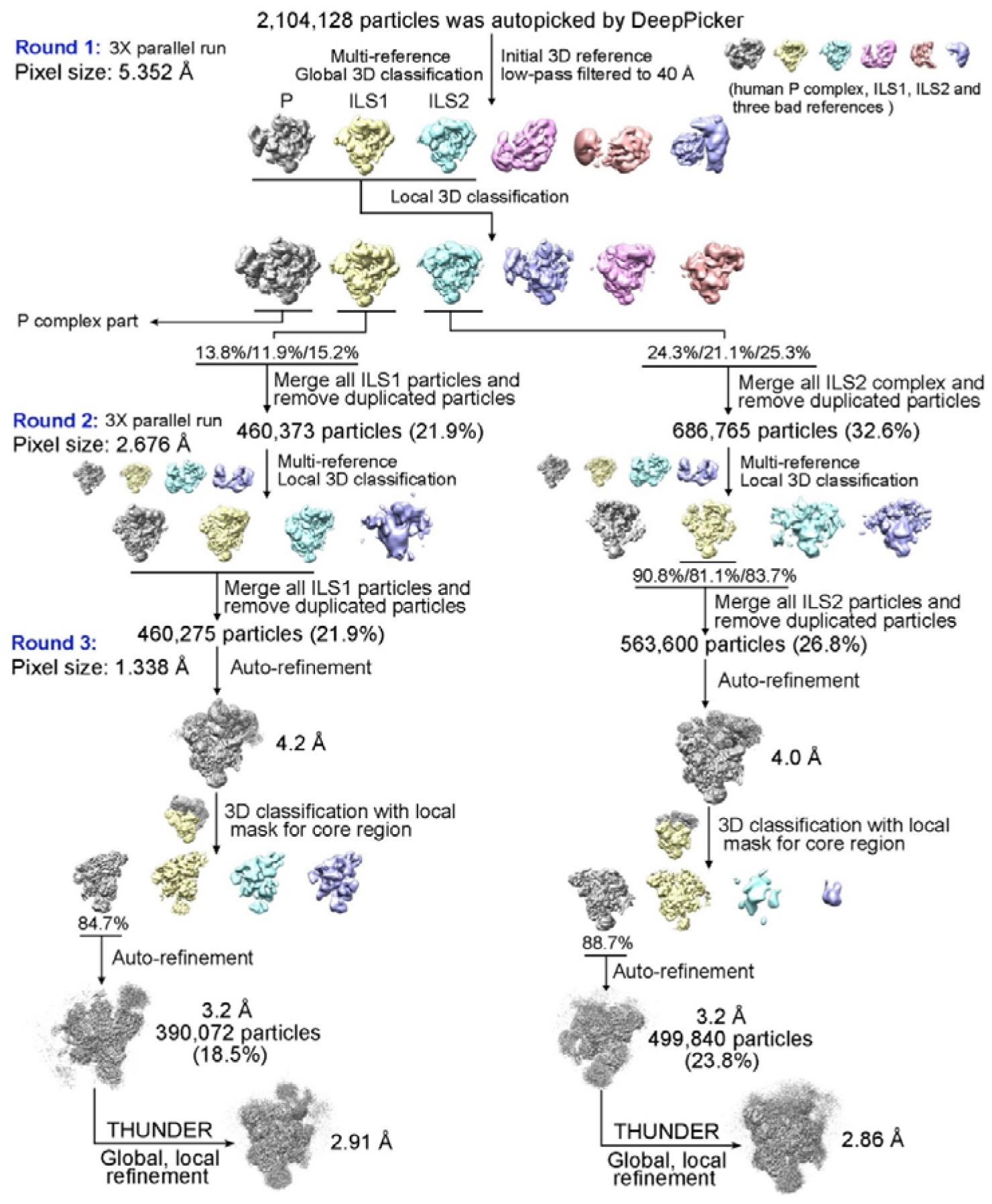
A flow chart description of the EM data processing and structure determination of the human spliceosomal ILS complexes. On the basis of the FSC value of 0.143, the final reconstruction has an average resolution of 2.91 Å for the ILS1 complex (prior to Prp43 recruitment) and 2.86 Å for the ILS2 complex (after Prp43 recruitment). Please refer to Methods for details.

**Extended Data Figure 4:**
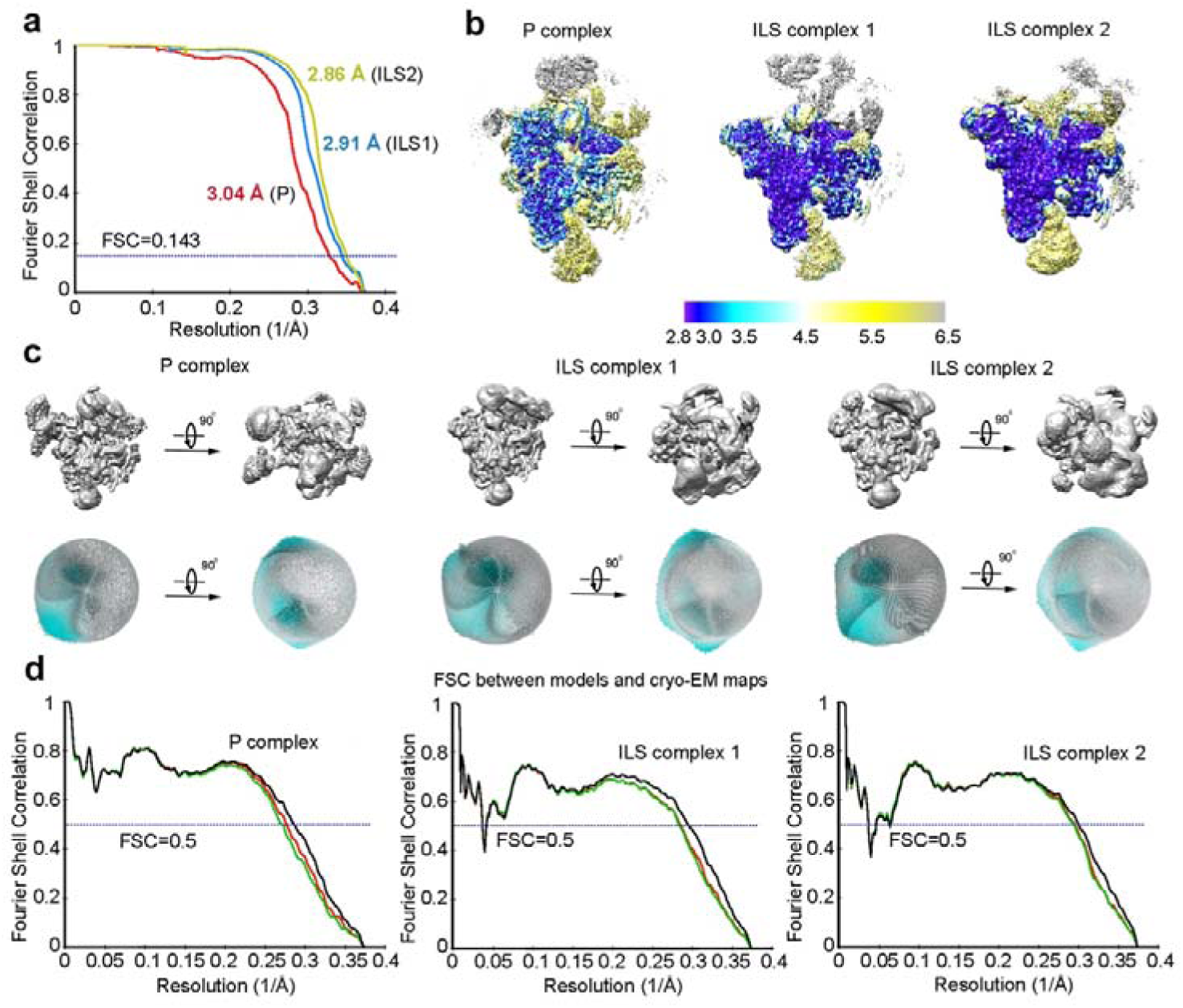
Cryo-EM analysis of the human spliceosomal P and ILS complexes. **a,** The average resolution is estimated to be 3.04 Å, 2.91 Å and 2.86 Å for the overall reconstruction of the P, ILS1, and ILS2 complexes, respectively, on the basis of the FSC criterion of 0.143. **b,** The local resolutions are color-coded for different regions of the P, ILS1, and ILS2 complexes. **c,** Angular distribution of the particles used for the reconstruction of the spliceosomal P, ILS1, and ILS2 complexes. Each cylinder represents one view and the height of the cylinder is proportional to the number of particles for that view. **d,** The FSC curves of the final refined model versus the overall map it was refined against (black); of the model refined in the first of the two independent maps used for the FSC calculation versus that same map (red); and of the model refined in the first of the two independent maps versus the second independent map (green). The generally similar appearances between the red and green curves indicates that the refinement of the atomic coordinates did not suffer from severe over-fitting.

**Extended Data Figure 5:**
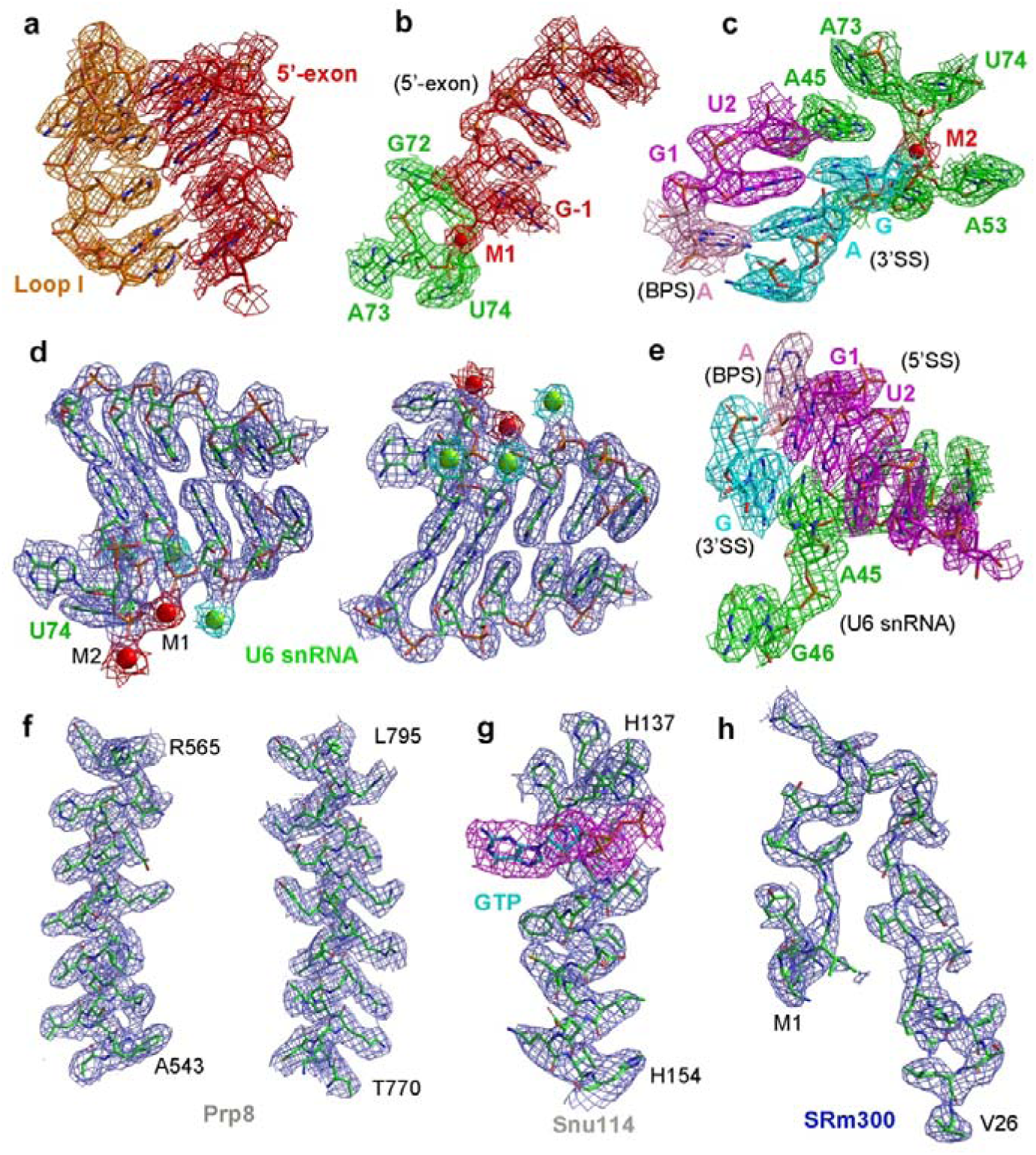
Representative EM density maps for select elements of the RNA and protein components in the human P complex. **a,** The EM density map for the duplex between 5’-exon and loop I of U5 snRNA. **b,** A close-up view on the EM density map surrounding the catalytic metal M1. **c,** A close-up view on the EM density map surrounding the 3’-splice site (3’SS). **d,** The EM density map for U6 snRNA, two catalytic metals (M1 and M2, colored red), and four structural metals (colored green). **e,** The EM density map of the duplex between 5’-splice site (5’SS) and U6 snRNA. **f,** Representative EM density maps of two helices from Prp8. **g,** A close-up view on the EM density map of GTP and a neighboring helix from Snu114. **h,** The EM density map of the splicing factor SRm300.

**Extended Data Figure 6:**
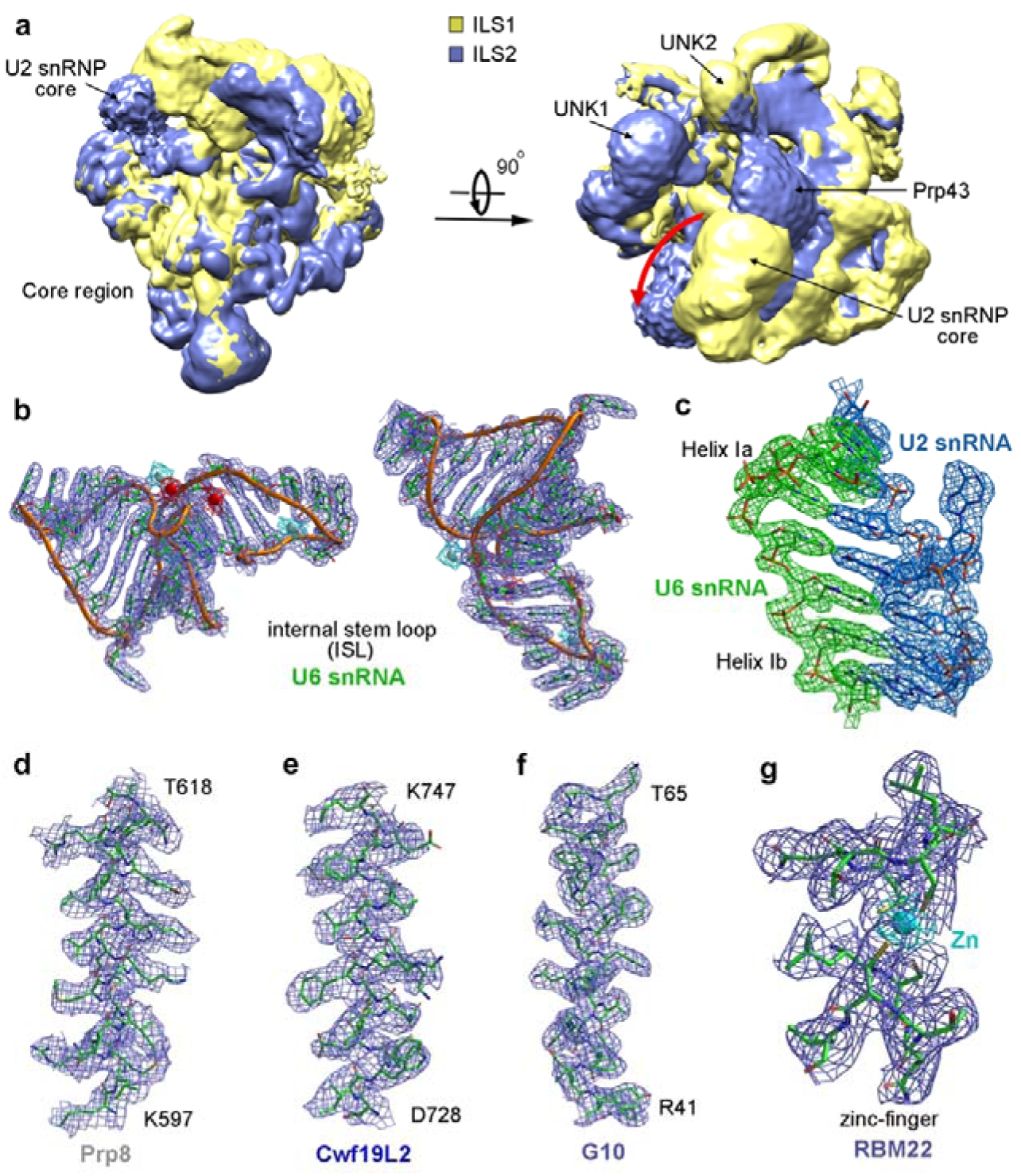
The EM density maps of the human ILS1 and ILS2 complexes. **a,** The overall EM density maps of the ILS1 and ILS2 complexes are low-pass filtered to 10 Å and overlaid together using Chimera. Two perpendicular views are shown. The core regions of the two ILS complexes are identical to each other. Compared to the ILS1 complex, the ILS2 complex has an extra lobe of density that is assigned to the ATPase/helicase Prp43. The U2 snRNP core has shifted during the ILS1-to-ILS2 transition. Each of the two ILS complexes contains two identical lumps of EM density near Prp43 that remain to be assigned. **b,** Two views of the EM density map for the internal stem loop (ISL) of U6 snRNA. **c,** A close-up view on the EM density map of helix I of the U2/U6 duplex. **d,** Representative EM density of one helix from Prp8. **e,** Representative EM density of one helix from Cwf19L2. **f,** Representative EM density of one helix from G10. **g,** A close-up view on the EM density of a zinc-finger from RBM22.

**Extended Data Figure 7:**
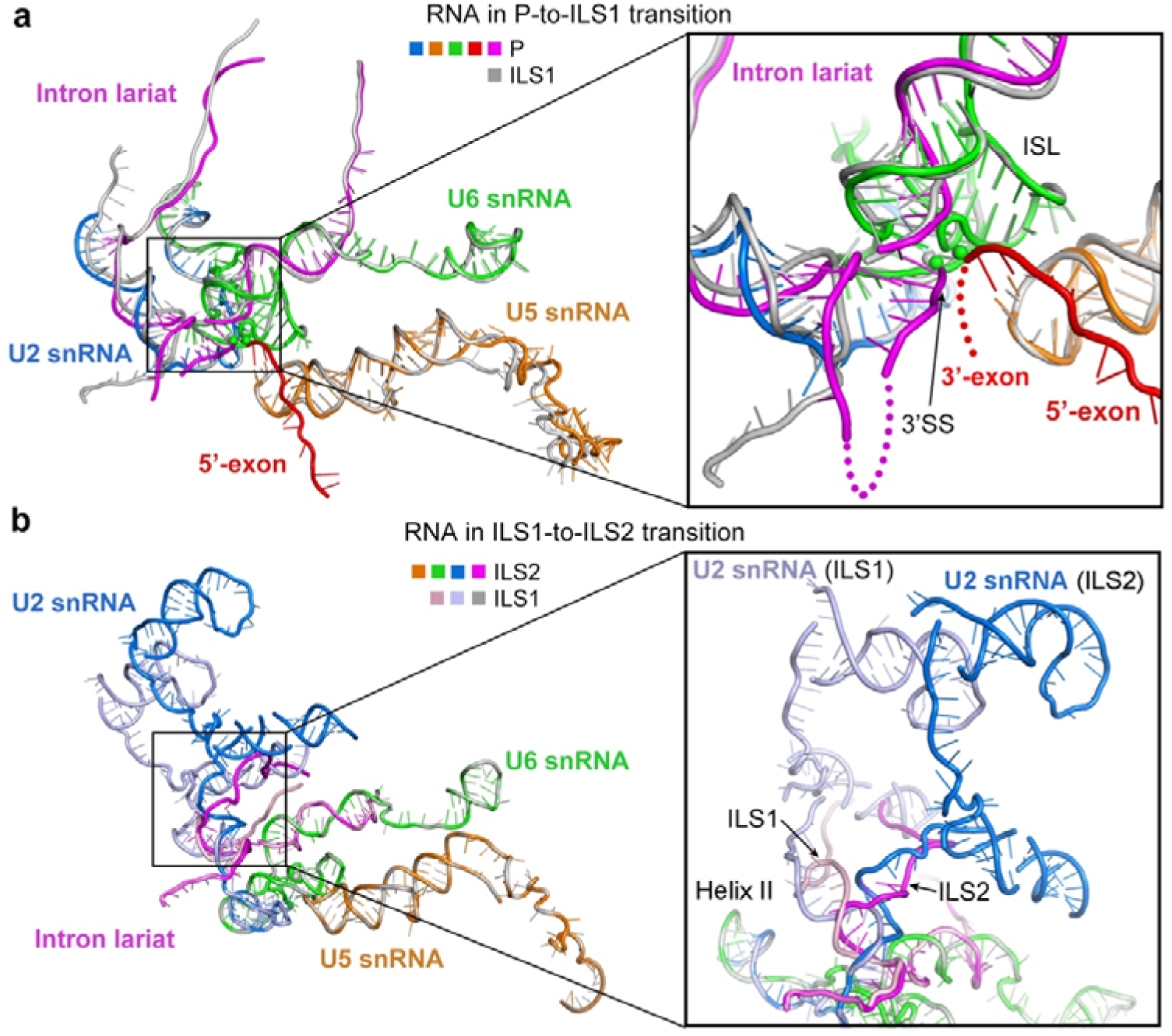
Comparison of the RNA elements and the splicing catalytic enter among the spliceosomal P, ILS1, and ILS2 complex. **a,** The RNA elements remain generally unchanged during the P-to-ILS1 transition, except that the ligated exon in the P complex has been dissociated in the ILS1 complex and the 3’-tail of the intron has rotated about 60 degrees away from the catalytic center. Shown here is the overlay of the overall RNA elements from the P and ILS1 complex (left panel) and a close-up view of the catalytic center (right panel). **b,** Rearrangements of the RNA elements during the ILS1-to-ILS2 transition. U5 and U6 snRNAs remain unchanged, whereas U2 snRNA is rotated about 40 degrees in the clockwise direction as shown. The U2/BPS duplex is also translocated.

**Extended Data Figure 8:**
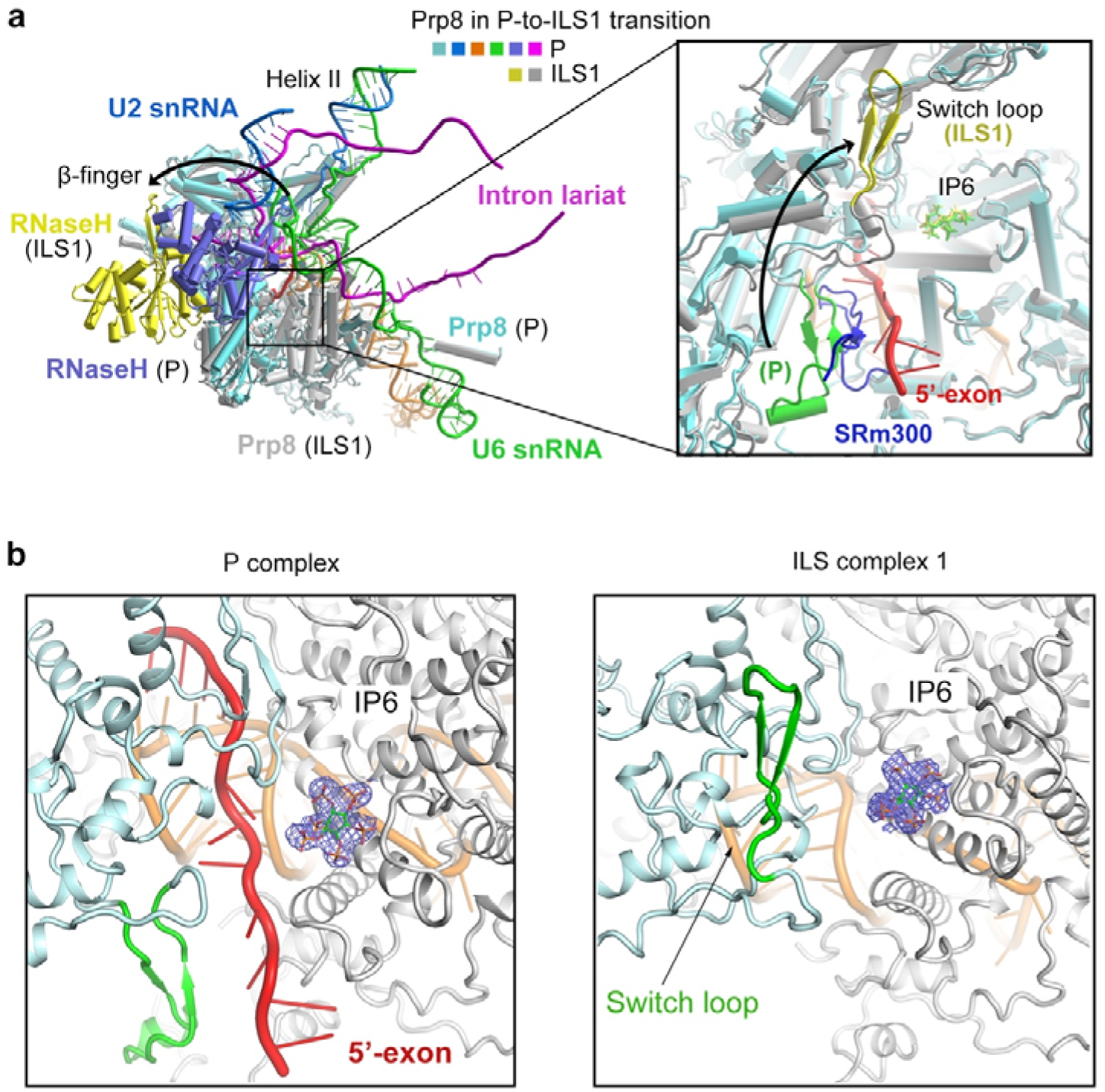
Conformational changes of Prp8 during the P-to-ILS1 transition. **a,** Superposition of Prp8 from the P and ILS1 complexes. The RNaseH-like domain from the P and ILS1 complex are colored blue and yellow, respectively. The N-domain and core from the P and ILS1 complexes are colored cyan and grey, respectively. The ß-finger is highlighted. In the right panel, a close-up view is shown on the Switch loop and a small molecule that has been tentatively assigned as inositol 1,2,3,4,5,6-hexaphosphate (IP6). **b,** Close-up views of IP6 from the P complex (left panel) and the ILS1 complex (right panel). The cryo-EM density maps for IP6 are shown.

**Extended Data Figure 9:**
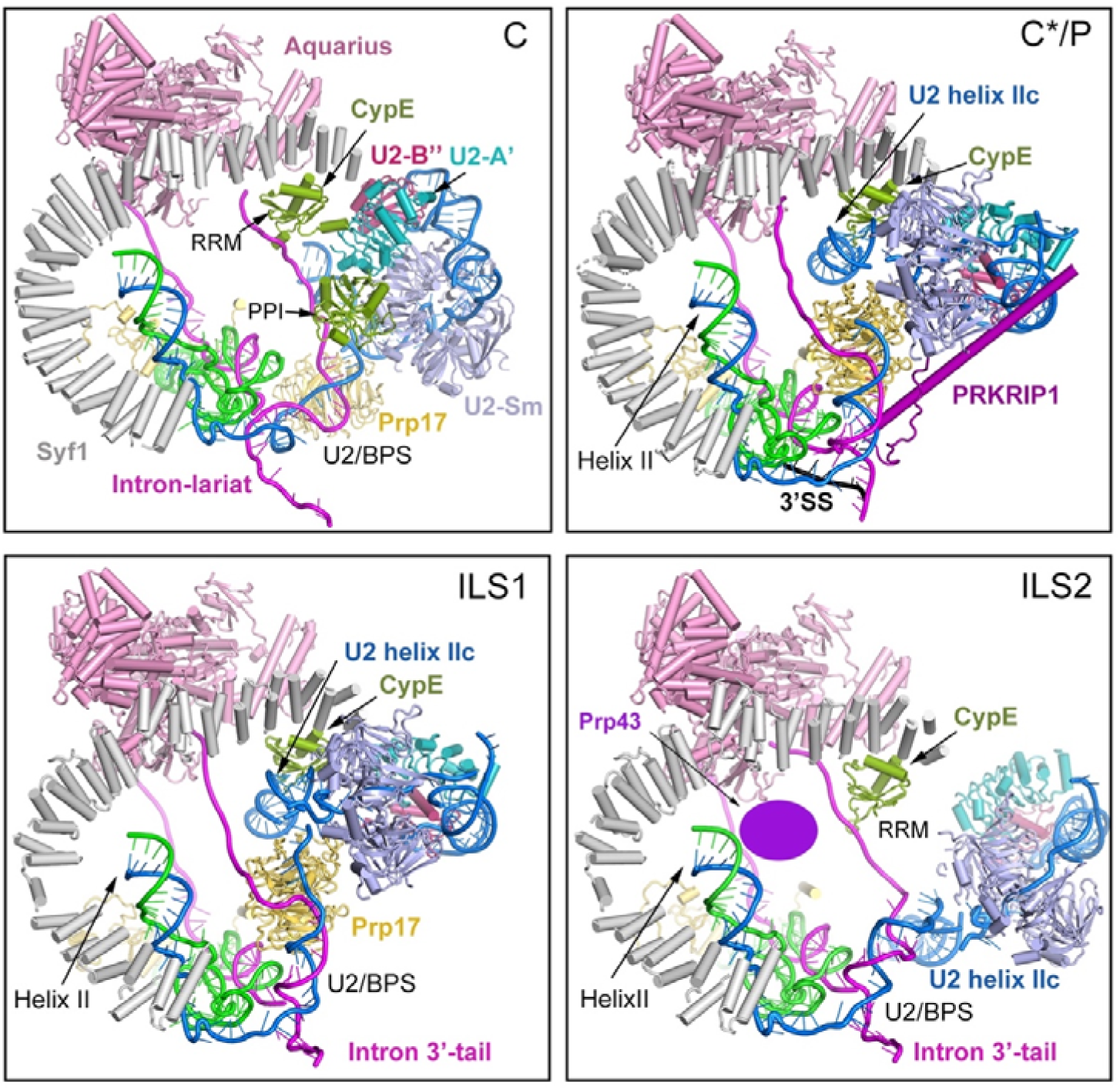
Movement of the U2 snRNP region during human spliceosome transition. U2 snRNP undergoes drastic translocation during remodeling of the human spliceosome. Shown here is the local conformations of the human spliceosomal C, C^*^/P, ILS1, and ILS2 complexes. In the C complex, the sequences upstream of the BPS are recognized by the RRM domain of CypE. Prp17 is yet to be loaded into the active site. During the C-to-C^*^ transition, U2 Sm ring is translocated by about 100 Å to the N-terminal region of Syf1, and the U2/BPS duplex is rotated away from active site. The step II factor Prp17 is now loaded between the U2/BPS duplex and the ISL. The intron sequences move away from CypE, which interacts with U2 helix IIc in the C^*^ complex. The splicing factor PRKRIP1 stabilizes the local conformation. In the P complex, the location of U2 snRNP remains unchanged. During the P-to-ILS1 transition, the local conformation remains unchanged except that PRKRIP1 is dissociated. In ILS2, Prp43 is recruited and located on the concave side of Syf1. The U2 snRNP core is translocated away from CypE, which is re-bound by the intron sequences upstream of the U2/BPS duplex. The WD40 domain of Prp17 is moved away from the active site and becomes disordered in ILS2 complex.

